# Diversification dynamics and (non-)parallel evolution along an ecological gradient in African cichlid fishes

**DOI:** 10.1101/2021.01.12.426414

**Authors:** A. A.-T. Weber, J. Rajkov, K. Smailus, B. Egger, W. Salzburger

## Abstract

Understanding the drivers and dynamics of diversification is a central topic in evolutionary biology. Here, we investigated the dynamics of diversification in the cichlid fish *Astatotilapia burtoni* that diverged along a lake-stream environmental gradient. Whole-genome and morphometric analyses revealed that divergent selection was essential at the early stages of diversification, but that periods in allopatry were likely involved towards the completion of speciation. While morphological differentiation was continuous, genomic differentiation was not, as shown by two clearly separated categories of genomic differentiation. Reproductive isolation increased along a continuum of genomic divergence, with a “grey zone” of speciation at ∼0.1% net nucleotide divergence. The quantification of the extent of (non-)parallelism in nine lake-stream population pairs from four cichlid species by means of multivariate analyses revealed one parallel axis of genomic and morphological differentiation among seven lake-stream systems. Finally, we found that parallelism was higher when ancestral lake populations were more similar.

## Introduction

The formation of new species – speciation – is a fundamental and omnipresent evolutionary process that has attracted much interest since Darwin’s seminal book from 1859 (*1*). Speciation is commonly defined as the evolution of reproductive isolation through the building up of barriers to gene flow (*2*). Speciation can occur ‘suddenly’ or gradually, along with the evolution of reproductive isolation between diverging populations (*2–4*). Sudden speciation is possible, for example, via hybridisation (*5*), polyploidization (*6*), or when a new mutation (e.g. a chromosomal inversion) directly leads to reproductive isolation (*7*). Typically, however, speciation has been considered a continuous process, during which genetic and phenotypic differences accumulate gradually between diverging populations and reproductive barriers become stronger until complete reproductive isolation is reached – a progression often referred to as speciation continuum (*4*). In the genic view of speciation, a small set of genes under divergent natural (or sexual) selection becomes resistant to gene flow at the initial stages of diversification, creating ‘genomic islands’ of strong differentiation; as the populations diverge, genetic differentiation expands across the genome, leading to stronger and more genome-wide patterns of differentiation (*4, 8, 9*). Speciation is considered “ecological” when barriers to gene flow arise as a result of ecologically-driven divergent selection (*10*). However, there are many empirical cases in which divergent selection initiates differentiation via local adaptation but is apparently not sufficient to complete speciation (e.g. *11* and references therein). This suggests that the factors driving initial population divergence may not be the same as those that complete speciation (*12*).

The dynamics of genomic differentiation during population divergence has been explored using both modelling (*3*) and empirical data (*13–15*). Simulations under a model of primary divergence with gene flow and divergent selection driving differentiation revealed that there can be a ‘sudden’ transition from a state of well-intermixed populations to two reproductively isolated entities (*3*), which has recently received empirical support (*13*). On the other hand, genomic divergence appears more gradual in other empirical systems (*14, 15*). So far, however, only few biological systems have been established that allow to jointly investigate early and late stages of differentiation by examining the dynamics of genomic and morphological differentiation along the entire speciation continuum.

Another outstanding question in evolutionary biology is to infer to what extent evolution is predictable. As biologists cannot ‘replay the tape of life’ (*16*), a common way to test the deterministic nature of selection is to examine evolutionary parallelism at phenotypic and genotypic levels across population pairs diverging along similar environmental gradients (*17*). For instance, parallel and non-parallel components in adaptive divergence have been reported in many fish species (see (*18*) for a review). A common observation arising from such studies is that, while some traits or genes indeed evolve in parallel, others in different systems do not. Furthermore, it has been reported that levels of parallelism tend to be lower in distantly related populations (*17*). However, this has rarely been formally tested in nature. Recently, the concept of (non-)parallel evolution has been introduced as a gradient ranging from truly parallel to completely divergent evolution instead of applying a binary classification (*19*). This allows to better quantify the extent of parallelism and to distinguish between convergence and parallelism (*19*). Specifically, the difference between convergent evolution and evolutionary parallelism is that in convergent evolution similar phenotypes or genotypes evolve from different initial conditions, whereas in parallel evolution initial conditions are similar (*19*).

East African cichlid fishes are important model taxa in speciation research, and constitute textbook examples of adaptive radiation characterized by rapid speciation accompanied by the evolution of substantial phenotypic, behavioural and ecological diversity (*20–22*). Here, we investigate the dynamics of genomic and morphological diversification along an environmental gradient in the East African cichlid fish *Astatotilapia burtoni* (Günther 1893) and across its distribution range. *Astatotilapia burtoni,* which occurs in African Lake Tanganyika (LT) and affluent rivers (Fig. 1A), was among the first five cichlid species to have their genomes sequenced (*23*). It is a generalist able to feed on a variety of food sources and can thrive in varied environments such as rivers and lakes (*24*), thus representing an ideal model species to investigate the dynamics of diversification along a lake-stream environmental gradient and across different geographic scales. It has previously been established that many tributaries of LT – be they small creeks or larger rivers – are inhabited by *A. burtoni* populations derived from lake fish, thereby forming ‘population pairs’ consisting of a source (that is, ancestral) population in the lake and a phenotypically distinct river population featuring habitat-specific adaptations in morphology and ecology (*24, 25*). For example, stream fish have a more inferior mouth position and a more slender body than lake fish (*24*). The different population pairs display varying levels of genomic differentiation, ranging from virtually no divergence to highly differentiated populations, and show strong signals of isolation-by-distance (*25*), highlighting that both genetic drift and divergent selection are at play in *A. burtoni* lake-stream divergence. Finally, a comprehensive phylogeographic study of *A. burtoni* across LT revealed that the populations from the North and the South of LT are genetically clearly distinct (*26*).

**Fig. 1.**
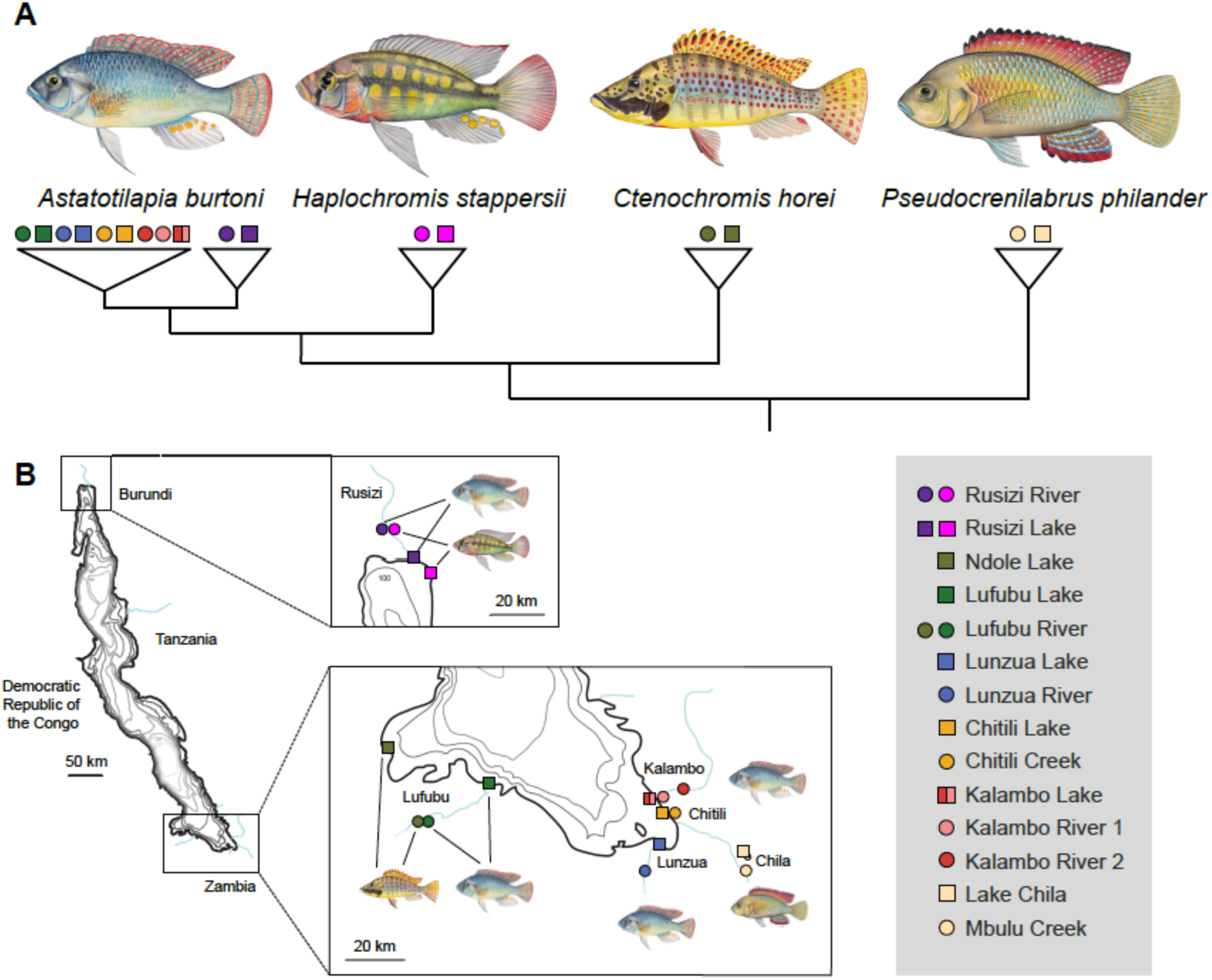
The study system comprising nine lake-stream population pairs in four cichlid fish species. (**A**) Illustrations of the four species used in this study and a schematic representation of their phylogenetic relationships (see Fig. S1 and ref.(*22*)). (**B**) Map of sampling localities and names of the different lake-stream population pairs, that is “systems”. *A. burtoni* (*N*_genomes_ = 132; *N*_morphometrics_ = 289), *H. stappersii* (*N*genomes = 24; *N*_morphometrics_ = 81)*, C. horei* (*N*_genomes_ = 24; *N*_morphometrics_ = 67) and *P. philander* (*N*_genomes_ = 24; *N*_morphometrics_ = 31).

To extend the comparative framework of this study beyond a single taxonomic unit, we also included lake-stream population pairs of three additional cichlid species belonging to the Haplochromini/Tropheini clade (*22*), namely *Haplochromis stappersii*, *Ctenochromis horei,* and *Pseudocrenilabrus philander* (Fig. 1A). Remarkably, two of these species co-occur in sympatry with *A. burtoni* in two of the largest tributaries to LT (*H. stappersii* in the Rusizi River in the North and *C. horei* in the Lufubu River in the South), providing an unprecedented opportunity to examine two replicates of lake-stream divergence and the extent of convergent evolution at genomic and morphological levels.

We used whole-genome resequencing, geometric morphometric analyses and mate-choice experiments to (*i*) assess the dynamics of genomic and morphological differentiation along the lake-stream environmental gradient and across geography in *A. burtoni*; (*ii*) evaluate to what extent genome-wide differentiation scales with reproductive isolation in *A. burtoni*; (*iii*) quantify genomic and morphological (non-)parallelism and convergence among nine lake-stream populations pairs from four haplochromine species; and (*iv*) test if levels of (non-)parallelism are higher when ancestral populations are more similar.

## Results

We performed whole-genome resequencing of 204 specimens of haplochromine cichlid fishes from LT and its surroundings (132 *A. burtoni* and 24 of each of the three additional haplochromine species) representing 17 populations. We included six *A. burtoni* lake-stream population pairs and one population pair for three additional species (*C. horei*, *H. stappersii,* and *P. philander*), totalling nine lake-stream population pairs (Fig. 1, Fig. S1, Tables S1 and S2; see Methods). Each population pair consisted of one lake and one stream population, whereby the lake population was sampled from a lake habitat close to the stream’s estuary. Each population pair, or ‘system’, was named after the respective river, except for the Lake Chila system (Fig. 1B). Note that from the Kalambo drainage we sampled two ecologically distinct river populations – one from a locality near the estuary where the river is deep and flows slowly (Kalambo 1) and the other one from ∼6 km upstream in a white-water environment (Kalambo 2) – resulting in two population pairs for this system (*24*). In addition, we quantified body shape of 468 specimens covering all 17 populations (Table S2). Finally, we evaluated the degree of reproductive isolation between *A. burtoni* populations displaying increasing levels of genomic divergence. To do so, we revisited published studies that examined the same *A. burtoni* populations as in the present study, and, in addition, we performed mate-choice experiments in the laboratory between the genetically most distinct *A. burtoni* populations from the North and the South of LT.

### The dynamics of genomic and morphological diversification

We first compared genomic (genome-wide *F_ST_*) and morphological (Mahalanobis distance, *D_M_*) differentiation across all *A. burtoni* populations, including the divergent populations from the North of LT, to examine the respective roles of environmental variation and geography in diversification. We calculated all pairwise *F_ST_* and *D_M_* comparisons between the 11 *A. burtoni* populations, resulting in 59 comparisons. We classified these comparisons according to environmental contrasts (lake-lake; lake-stream; stream-stream) and to the geographic distance between populations (1-40 km: South-East of LT; 70-140 km: East-West of South LT; 700-750 km: North-South of LT; see Fig. 1B). We found that morphological differentiation was gradual, with *D_M_* values ranging from 2.1 (Lunzua lake versus Kalambo lake) to 7.6 (Rusizi lake versus Lufubu River) (Fig. 2A, Table S3). In all three geographic groups, the most similar system-pairs were lake-lake comparisons (i.e. similar environments), whereas the most different ones were lake-stream comparisons, suggesting that ecological factors impact body shape differentiation. On the other hand, a wide range of *D_M_* values was observed within a small geographic range (1-40 km: Lunzua lake versus Kalambo lake: *D_M_* = 2.1; Chitili River versus Kalambo lake: *D_M_* = 6.3), suggesting that body shape differentiation can be rapid across small geographic distances and low levels of genome-wide divergence (Fig. 2A, Fig. S2).

**Fig. 2.**
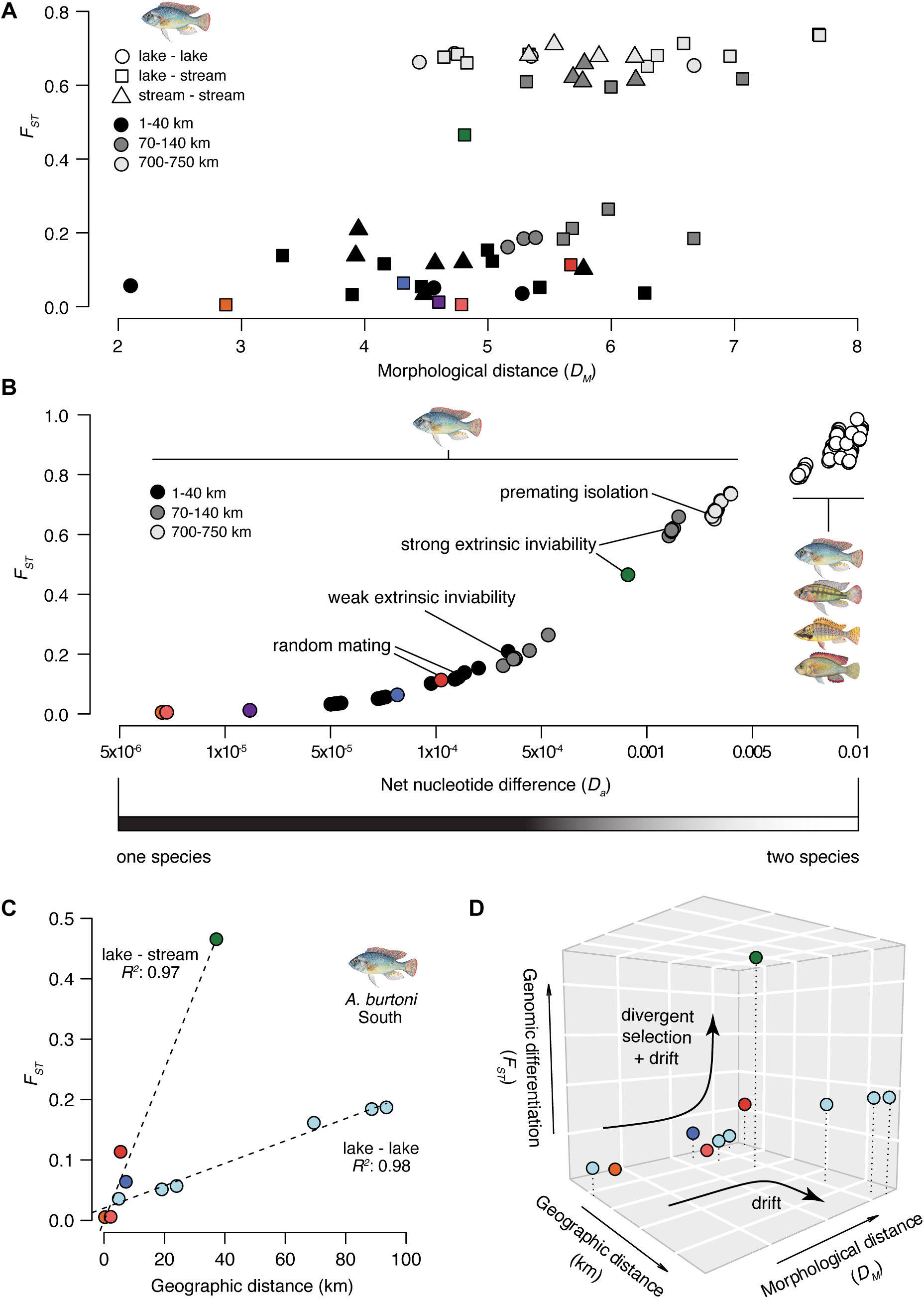
Dynamics of diversification in *Astatotilapia burtoni*. **(A)** Genetic distances (genome-wide *F_ST_*) between all *A. burtoni* populations are plotted against morphological distances (Mahalanobis distances (*D_M_*) calculated from body shape). The geographic distance between populations and the habitat type for each comparison are highlighted with symbol colour (light grey, dark grey, black) and shape (circle, square, triangle), respectively. The sympatric lake-stream systems are highlighted in colours (same colour coding as in Fig. 1). Morphological distances increase gradually whereas two groups of genetic distances are observed; the ‘one-species’ (*F_ST_* < 0.3) and the ‘two-species’ category (*F_ST_* > 0.6). The Lufubu lake-stream system (green) is the only intermediate comparison (distance 40 km). Comparisons within the ‘one-species’ category were all found at short geographic distances (1-40 km), whereas at intermediate geographic distances (70-140 km), comparisons from both categories could be found. Finally, at large geographic distances (700-750 km), all comparisons belong to the ‘two-species’ category, irrespective of the environment. (**B**) Genome-wide *F_ST_* plotted against the net nucleotide difference (*D_a_* = *d_XY_* –(π_1_+π_2_)/2; a proxy for time since divergence - note the log-scale for the x axis) in 55 pairwise comparisons of all *A. burtoni* populations, representing the speciation continuum. Pairwise comparisons among species (*P. philander, C. horei, H. stappersii*) are reported for comparative purposes. Genomic differentiation accumulates fast during early stages of divergence (*F_ST_* < 0.3; that is, the ‘one-species’ category) but then slows down as *D_a_* increases (*F_ST_* > 0.6; that is, the ‘two-species’ category). The sympatric population pair from the Lufubu system is intermediate (*F_ST_ A. burtoni*: 0.46). Levels of reproductive isolation increase along a continuum of genomic differentiation (see Table 1 for details on the experiments and population used). (**C**) Isolation-by-distance in the two comparison categories ‘lake-stream’ (The colour coding for lake-stream systems is the same as in Fig. 1) and ‘lake-lake’ (light blue) in the southern populations of *A. burtoni* (that is, within the one-species category). R^2^: Pearson’s correlation coefficient. (**D**) Trajectories for three differentiation axes: morphology, geography and genetics. In ‘lake-stream’ comparisons, morphological differentiation builds up first, then genomic differentiation increases sharply, likely due to the combined effect of divergent selection and drift at small geographic distance. In contrast, in the absence of strong divergent selection (that is, in the ‘lake-lake’ comparisons), morphological differentiation builds up first, then genomic differentiation accumulates only moderately, which is likely due to non-adaptive processes such as genetic drift.

**Table 1:** Summary of experimental testing of reproductive isolation in *A. burtoni*.

| Populations | $F_{ST}$ | Genetic markers | Experimental set-up | Main results | Reproductive isolation | Reference |
| --- | --- | --- | --- | --- | --- | --- |
| Kalambo lake × Kalambo River 2 | 0.113 | Whole-genome | Transplant experiment in the lake environment | Adaptive phenotypic plasticity | Weak for wild-caught adults; not observed for F1 | (27) |
| Kalambo lake × Lunzua River | 0.123 | Whole-genome | Common garden experiment in mesocosms | Random mating | Not observed | (24) |
| Kalambo lake × Ndole lake | 0.189<sup>1</sup> | Whole-genome<sup>1</sup> | Common garden experiment in mesocosms | Immigrant and extrinsic hybrid inviability (weak) | Extrinsic prezygotic and postzygotic isolation (weak) | (28) |
| Ndole lake × Lufubu River | 0.472<sup>1</sup> | Whole-genome<sup>1</sup> | Common garden experiment in mesocosms | Immigrant and extrinsic hybrid inviability (strong) | Extrinsic prezygotic and postzygotic isolation (strong) | (28) |
| Kalambo lake × Lufubu River | 0.624 | Whole-genome | Common garden experiment in mesocosms | Immigrant and extrinsic hybrid inviability (strong) | Extrinsic prezygotic and postzygotic isolation (strong) | (28) |
| Kalambo lake × Rusizi lake | 0.693 | Whole-genome | Laboratory mate-choice experiment | Partial assortative mating | Premating isolation | This study |

<sup>1</sup> Genetic distances inferred from the Lufubu lake population rather than Ndole lake population (both populations are geographically close and belong to the same genetic cluster (24)).

Contrary to morphological differentiation, we observed a sudden increase in genomic differentiation, with two clearly separated categories in *F_ST_*-values (≤ 0.3 and ≥ 0.6), hereafter referred to as the ‘one-species category’ and the ‘two-species category’, respectively (Fig. 2A). The comparisons in the ‘one-species category’ included all south-eastern populations and the Lufubu lake population, while comparisons in the ‘two-species category’ included all south-eastern populations versus the Lufubu River population, and all southern populations versus both northern populations. Geographic distance did not appear to be the main driver of this separation, as comparisons including moderate geographic distances between populations (70-140 km) were found in both *F_ST_* categories. Rather, it appears that the separation at moderate geographic distance was driven by ecological factors, as the comparisons in the ‘two-species category’ included all south-eastern populations versus the Lufubu River population. Interestingly, there was one comparison with intermediate *F_ST_*-value between the one-species and the two-species categories, namely the population pair from the Lufubu system (Lufubu lake versus Lufubu River: *F_ST_* = 0.46). Demographic modelling of this population pair indicated that the most likely scenario of divergence between these populations included a past secondary contact event, that is, one or several periods of allopatric divergence in the past with current ongoing gene flow (Fig. S3; Table S4).

Next, we investigated the increase in genomic differentiation against a proxy of time since population divergence, net nucleotide difference (*D_a_*). To extend the comparative framework encompassing a whole continuum of genomic divergence from populations to species, we here included the between-species comparisons of the three additional haplochromine species *H. stappersii, C. horei* and *P. philander*. We found a rapid increase in *F_ST_* at low levels of *D_a_*, which slowed down as *D_a_* increased further. Given this trend, we fitted a logarithmic model to the data which was highly significant (linear regression *F_ST_* ∼ log(*D_a_*): *P* < 2.2e^-16^; Pearson correlation coefficient: *R^2^* = 0.96). Therefore, to better visualize the dynamics of diversification at early stages of genomic differentiation, *D_a_* was plotted on a logarithmic scale (Fig. 2B). The one-species category encompassed *D_a_* values ranging from 5x10^-6^ to 5x10^-4^, while the two-species category features *D_a_* values of 0.002 and above. With *D_a_* ∼0.001, the population pair from the Lufubu system occupied the ‘grey zone’ of speciation between the one-species and two-species categories.

### Reproductive isolation begins establishing at low levels of genome-wide differentiation

As a next step, we assessed to what extent the observed levels of genome-wide differentiation scale with the degree of reproductive isolation between populations in *A. burtoni*. To this end, we revisited available data from previous experiments that used the same *A. burtoni* populations as in the present study (*24, 27, 28*) and conducted additional mate-choice experiments in the laboratory between the genetically most distinct populations of *A. burtoni* from the North and the South of LT (*26*) (Table 1, Fig. S4).

Previous experiments involving populations that feature low levels of genomic divergence (Kalambo lake versus Kalambo River 2: *F_ST_* = 0.11 (*27*); Kalambo lake versus Lunzua River: *F_ST_* = 0.12 (*24*)) revealed random mating patterns with respect to source population, suggesting a lack of reproductive isolation. In a mesocosm experiment with populations at a slightly higher level of genome-wide differentiation, yet still within the one-species category (Kalambo lake versus Ndole lake: *F_ST_* = 0.18), weak levels of extrinsic hybrid inviability were found (*28*). In contrast, populations at intermediate (Ndole lake versus Lufubu River: *F_ST_* = 0.47) and high (Kalambo lake versus Lufubu River: *F_ST_* = 0.62) levels of genome-wide divergence showed strong levels of extrinsic hybrid inviability in the same mesocosm experiment (*28*). Therefore, evidence of reproductive isolation was detected for populations in the ‘grey zone’ of speciation and in the two-species category, indicating that our species categories that are solely based on levels of genome-wide differentiation in *A. burtoni* are biologically meaningful. In support of this, our new laboratory-based mate-choice experiments targeting two populations that feature one of the highest genome-wide *F_ST_*-values (Kalambo lake versus Rusizi lake: *F_ST_* = 0.69) revealed signatures of assortative mating with respect to source population – at least in a multi-sensory laboratory setting allowing for a combination of mating cues (Fig. S4 and Appendix 1). This suggests that premating reproductive isolation mechanisms are at play between the genetically most distinct *A. burtoni* clades – the northern and southern lineages (*26*) – at a level of genomic differentiation that is similar to the one typically observed *between* other haplochromine species (Fig. 2B).

### Divergent selection and drift accelerate genome-wide differentiation

We then investigated the relative influence of divergent selection and geography on genomic and morphological diversification in *A. burtoni*. To do so, we compared the more closely-related southern populations of *A. burtoni* focusing on within habitat (lake-lake) versus between habitats (lake-stream) comparisons. The latter consisted of the five *A. burtoni* lake-stream systems from the South of LT (i.e. Chitili; Kalambo 1; Lunzua; Kalambo 2; Lufubu) (Fig. 1B), whereas the lake-lake comparisons consisted of all pairwise comparisons of lake populations from that area. In both cases, we found an increase in *F_ST_* over geographic distance (Fig. 2C), which is compatible with an isolation-by-distance scenario for lake-stream (linear regression: *P* = 0.0013; Pearson correlation coefficient: *R^2^* = 0.97) and lake-lake (Mantel test: *P* = 0.08; Pearson correlation coefficient: *R^2^* = 0.98) contrasts. Interestingly, genomic differentiation increased much stronger with respect to geographic distance when populations were compared that inhabit contrasting environments (lake-stream population comparisons; i.e. in the presence of divergent selection and drift) than when they inhabit similar environments (lake-lake comparisons; i.e. primarily in the presence of drift), suggesting that divergent selection outweighs isolation-by-distance in diversification in *A. burtoni* (Fig. 2C).

Demographic modelling within lake-stream population pairs suggested that there is ongoing gene flow between lake and stream fish in all population pairs (Fig. S3; Table S4) and that the effective population size of most stream populations was smaller than the size of the respective lake population (Table S4), which is compatible with the scenario that the respective rivers were colonized from lake stocks (*24, 25*). As the impact of genetic drift is stronger in small populations, drift may also have contributed to the increased genomic divergence of stream populations. Interestingly, the diversification trajectories were similar between lake-stream and lake-lake comparisons at low levels of genetic, morphological and geographic distances (Fig. 2D), but diverged as genomic differentiation built up much more rapidly compared to geographic distance in the presence of divergent selection and increased drift (lake-stream comparisons) (Fig. 2D). This corroborates the notion that the environment (via divergent selection) and demographic events play a crucial role in differentiation trajectories in *A. burtoni* at early stages of differentiation.

### Little overlap between differentiation regions among independent lake-stream systems

We then turned our attention to the question of parallel genomic and morphological evolution along the lake-stream environmental gradient. To do so, we compared the six *A. burtoni* lake-stream population pairs and used three additional population pairs from different haplochromine species to extend the comparative framework to between-species comparisons. Notably*, H. stappersii* and *C. horei* are found in sympatry with *A. burtoni* in the Rusizi River and in the Lufubu River, respectively, providing an unprecedented opportunity to examine two replicates of lake-stream colonization across species. The lake-stream population pairs displayed contrasting levels of genome-wide differentiation, ranging from *F_ST_* = 0.005 (Chitili) to *F_ST_* = 0.465 (Lufubu) in *A. burtoni* (Fig. 3). Differentiation between lake-stream population pairs in the other three haplochromines ranged from low (*H. stappersii*; *F_ST_* = 0.046) to intermediate (*C. horei*; *F_ST_* = 0.532) to high (*P. philander*; *F_ST_* = 0.733), corroborating that the *P. philander* populations may actually represent two distinct species, as suggested by their different sex determining systems (*29*). Consistent with this, the *P. philander* populations display relatively high levels of absolute divergence (*d_XY_* = 3.6x10^-3^), which are above the levels of divergence measured between the northern and southern *A. burtoni* lineages (*d_XY_* = 3.0-3.3x10^-3^) (Table 2).

**Fig. 3.**
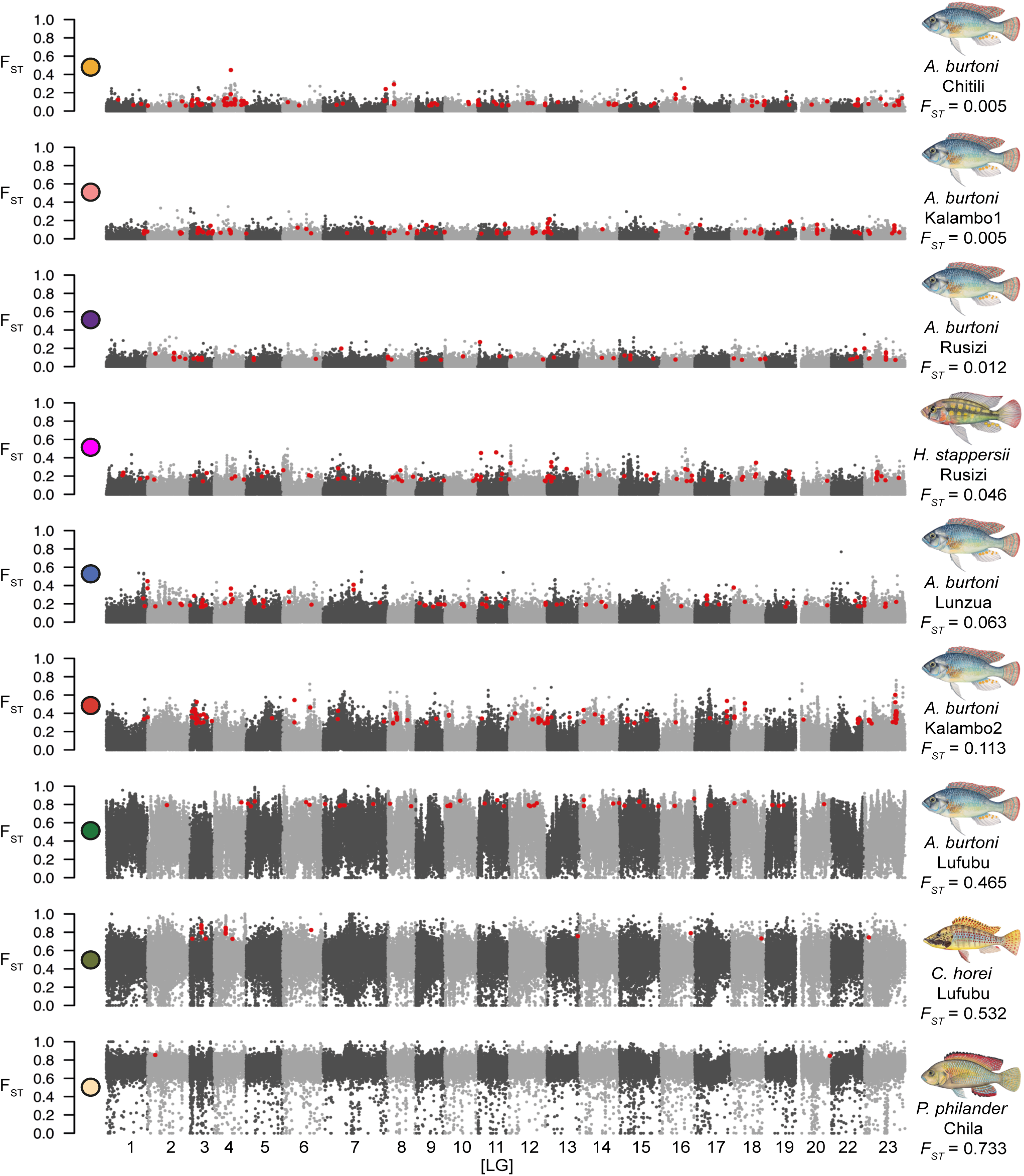
Distribution of genome-wide *F_ST_* for the nine lake-stream systems sorted by increasing genome-wide *F_ST_*-value. Each dot represents an *F_ST_*-value calculated in a 10-kb window along each linkage group. Linkage groups are highlighted in different shades of grey. Regions of differentiation (overlap of the top 5% values of *F_ST_*, *d_XY_* and *π*) are highlighted in red.

**Table 2:** *d_XY_* values between divergent populations and species used in this study. *d_XY_* values are presented as a 10^-3^ factor for readability. *A. burtoni* ‘South’ include all populations from south-east of Lake Tanganyika (LT) including Lufubu Lake population. *A. burtoni* ‘Lufubu’ includes the Lufubu River population. *A. burtoni* ‘North’ includes both populations (Rusizi lake and Rusizi River) from the North of LT.

|  | *A. burtoni*<br>‘South’ | *A. burtoni*<br>‘Lufubu’ | *A. burtoni*<br>‘North’ | *H. stappersii* | *C. horei* | *P. philander* |
| --- | --- | --- | --- | --- | --- | --- |
| *A. burtoni*<br>‘South’ | 0.9-1.3 |  |  |  |  |  |
| *A. burtoni*<br>‘Lufubu’ | 1.7-2.1 | 0 |  |  |  |  |
| *A. burtoni*<br>‘North’ | 3.0-3.2 | 3.3 | 1.0 |  |  |  |
| *H. stappersii* | 6.7-6.8 | 6.7 | 6.4-6.5 | 1.6 |  |  |
| *C. horei* | 8.4-8.5 | 8.4-8.5 | 8.4-8.5 | 8.5-8.7 | 1.1 |  |
| *P. philander* | 10.5-10.9 | 10.5-10.8 | 10.6-10.9 | 10.7-11.0 | 9.6-10.0 | 3.6 |

We next investigated genomic regions of high differentiation between lake-stream population pairs and evaluated to which extent such outlier regions (defined here as the intersection between the top 5% 10-kb windows with respect to *F_ST_*, *d_XY_*, and absolute value of *π* difference) are shared between population pairs and species. Our analyses revealed between 2 and 101 outlier regions of high differentiation per lake-stream population pair and that these regions were between 10-70 kb in length (red dots in Fig. 3, Table S5). It has been shown that heterogeneity in crossover rates can produce contrasting patterns of genomic differentiation between diverging populations that are not due to divergent selection. These are manifested, for example, in greater levels of differentiation near chromosome centres (where crossover rates are low) compared to the peripheries (where crossover rates are high) (*30*). In our case, we did not find evidence for an accumulation of differentiation regions in the chromosome centres in any of the lake-stream population pairs nor when all 525 outlier regions were considered jointly (Table S5), suggesting that our results reflect true signatures of divergent selection.

We then evaluated to what extent differentiation regions were shared among lake-stream population pairs and species. We found that there was little overlap (Fig. 4A) and that no such region of high differentiation was shared between more than two systems (Fig. 4A). The regions of high differentiation were distributed across all linkage groups, although there seems to be an overrepresentation on LG3 (64 out of 525), which remained when correcting for chromosome length (Table S5). The 525 differentiation regions contained a total of 637 genes. However, there was no obvious overrepresentation in functional enrichment with respect to Gene Ontology categories. The only exception was the Lunzua lake-stream comparison, in which outlier genes were significantly enriched for the molecular function “binding” (Table S6). The 25 genes located in the 19 differentiation regions shared between two systems also showed no functional enrichment (Table S7).

**Fig. 4.**
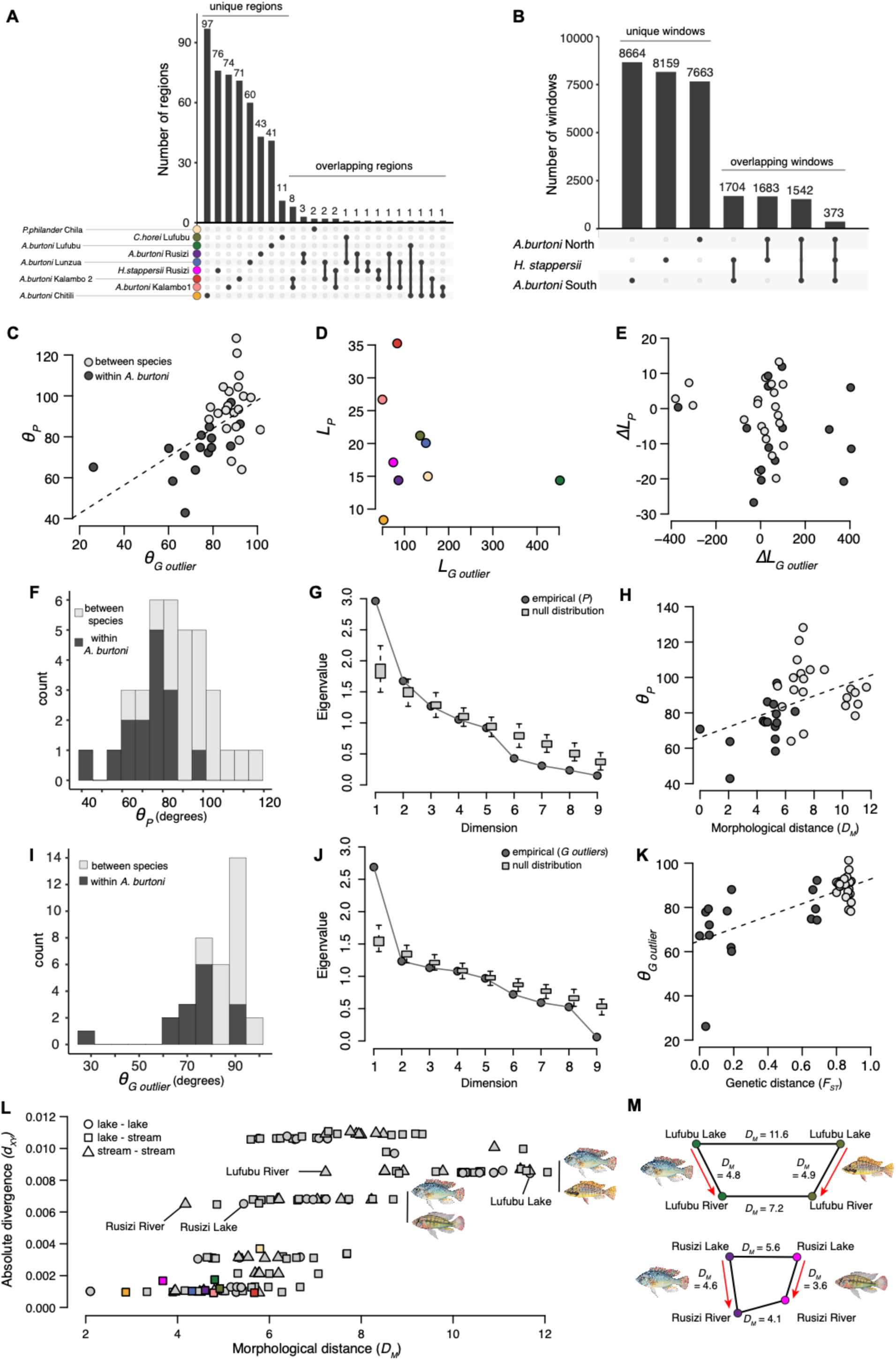
(Non-)parallel and convergent evolution among nine lake-stream cichlid population pairs. (**A**) Number of unique and overlapping differentiation regions (overlap of top 5% values of *F_ST_*, *d_XY_* and *π*) in the nine lake-stream population pairs. (**B**) Number of unique and overlapping differentiation windows in the three sets of ‘core’ outliers (core model of differentiation from BayPass) from *A. burtoni* northern populations, *A. burtoni* southern populations and *H. stappersii*. (**C**) The angles of phenotypic (*θ_P_*) and genetic outlier (*_θG outlier_*) lake-stream divergence vectors are positively correlated (Mantel test: *P* = 0.0075; Pearson correlation coefficient: R^2^=0.27). The dashed line indicates significant correlation at the 0.05 level. (**D**) The lengths of phenotypic (*L_P_*) and genetic outlier (*L_G outlier_*) lake-stream divergence vectors are not correlated (Linear regression model: *P*=0.55). The colour scheme is the same as in panel A. (**E**) The differences between phenotypic (ΔL_P_) and genetic outlier (ΔL_G_outlier_) vector length are not correlated (Mantel test: *P*=0.26). (**F**) Histogram of the 36 (pairwise) angles between lake-stream phenotypic divergence vectors (*θ_P_*) in degrees. Within *A. burtoni* and between species comparisons are highlighted in different shades of grey. (**G**) a multivariate analysis of phenotypic parallelism reveals one significant dimension of parallel evolution (the first empirical eigenvalue is higher than the null Wishart distribution). (**H**) The angles of phenotypic divergence vectors (*θ_P_*) and morphological distances (*D_M_*) between ancestral (i.e. lake) populations are positively correlated (Mantel test: *P* = 0.045; R^2^ = 0.17). For example, *θ_P_* between the Lunzua and Kalambo2 lake-stream systems is plotted against the Mahalanobis distance between Lunzua lake and Kalambo lake fish. In other words, if lake populations of two lake-stream systems are more similar morphologically, their direction of phenotypic divergence tends to be more parallel (have a small *θ_P_*). (**I**) Histogram of the 36 (pairwise) angles between lake-stream genetic outlier divergence vectors (*θ_G outlier_*) in degree. (**J**) a multivariate analysis of genetic parallelism reveals one significant dimension of parallel evolution (the first empirical eigenvalue is higher than the null Wishart distribution). (**K**) The angles of genetic outlier divergence vectors (*θ_G outlier_*) and genetic (*F_ST_*) distances between lake populations are positively correlated (Mantel test: *P*=0.0039; R^2^=0.46). (**L**) Absolute divergence (*d_XY_*) is plotted against morphological distance (*D_M_*) for 136 pairwise comparisons across all populations from the four species used in this study. A wide range of morphological distances can be observed for a same amount of genomic divergence. The sympatric populations (*A. burtoni* and *H. stappersii* from the Rusizi system; *A. burtoni* and *C. horei* from the Lufubu system) are highlighted. The habitat type for each comparison is highlighted with different symbols (circle, square, triangle). The colour scheme is the same as in panel A. (**M**) Smaller morphological distances between sympatric riverine populations compared to sympatric lake populations reveals body shape convergence in the riverine populations (*A. burtoni* and *H. stappersii* from the Rusizi system; *A. burtoni* and *C. horei* from the Lufubu system).

In situations of a shared evolutionary history of the populations in question, such as in our set-up, frequency-based outlier detection methods may not be the most appropriate way to detect regions under selection. Thus, we also performed Bayesian analyses of selection at the SNP level that take into account population relatedness (*31*). Due to their high levels of genome-wide differentiation, the population pairs of *C. horei* and *P. philander* were excluded from these analyses, as signatures of selection and drift cannot easily be disentangled in such cases. For the same reason, we treated the northern and southern populations of *A. burtoni* as separate units. We identified 1,704 shared 10-kb windows of differentiation between *A. burtoni* from the South and *H. stappersii*, 1,683 shared windows between *A. burtoni* from the North and *H. stappersii*, and 1,542 shared windows between *A. burtoni* from the North and from the South (Fig. 4B). In total, 373 windows were shared among the three core sets, containing 367 genes (Fig. 4B; Table S8). Interestingly, some genes involved in sensory perception (sound and light) were overrepresented in the common set of outliers between *H. stappersii* and the southern *A. burtoni* populations.

### (Non-)parallel evolution in lake-stream divergence

We further aimed to assess the extent of phenotypic and genomic (non-)parallelism among lake-stream pairs of haplochromine cichlids. To examine how (non-)parallel the nine lake-stream population pairs are, we performed vector analyses (*19, 32*) using the morphological (37 traits and landmarks representing body shape) and genomic (outlier and non-outlier SNPs) data at hand (outlier SNPs: BayPass outliers, based on 78 principal components from a genomic Principal Component Analysis (PCA), potentially impacted by natural selection; non-outlier SNPs: all SNPs but excluding BayPass outliers). Following Stuart et al. (*33*) we quantified variation in lake-stream divergence by calculating vectors of phenotypic and genomic differentiation for each lake-stream population pair, whereby the length of a vector (*L*) represents the magnitude of lake-stream divergence, and the angle between two vectors (*θ*) informs about the directionality of divergence. Accordingly, two lake-stream systems are more ‘parallel’ if *θ* is small (similar direction of divergence) and *ΔL* (difference in length between two vectors) is close to zero (similar magnitude of divergence) (*19*).

Most comparisons between independent lake-stream population pairs fell into the category ‘non-parallel’, with close to orthogonal (∼90°) angles of differentiation in both phenotype (*θ_P_*) and genotype (*θ_G_outlier_* and *θ_G_*) (Fig. 4F, I, Fig. S5D). However, parallelism was higher when only the within *A. burtoni* comparisons were considered, with values of *θ_P_* and *θ_G_outlier_* between 70° and 80° in many cases (Fig. 4F, I), but not for *θ_G_* (Fig. S5D). More clear signatures of parallelism were found in closely related *A. burtoni* lake-stream systems, with the Lunzua-Kalambo1 pair being the most ‘parallel’ system at the phenotype level (*θ_P_*: 42°) and the Chitili-Kalambo1 pair the most ‘parallel’ at the genotype level (*θ_G_outlier_*: 26°). Overall, the directions of phenotypic (*θ_P_*) and genetic (*θ_G_outlier_*) differentiations were significantly correlated (Mantel test: *P* = 0.0075; Pearson correlation coefficient: *R^2^* = 0.27; Fig. 4C), whereas their magnitudes were not (Mantel test *ΔL_P_* and *ΔL_G_outlier_*: *P* = 0.26; Linear regression model *L_P_* and *L_G_outlier_*: *P* = 0.55; Fig. 4D, E). In contrast, none of the non-outlier genetic vectors were correlated to phenotypic vectors (*θ_P_* versus *θ_G_* and *ΔL_P_* versus *ΔL_G_*: Mantel tests: *P* = 0.34; *P* = 0.35; *L_P_* and *L_G_*: Linear regression: *P* = 0.93; Fig. S5A-C).

It has recently been proposed to examine the vectors of divergence using a multivariate approach as a complementary set of analyses to investigate (non-)parallelism and convergence (*34*). For each dataset (phenotype, genotype outlier and genotype non-outlier), we thus calculated the eigen decomposition of the respective C matrix (*m* lake-stream systems x *n* traits). For the phenotype data, we found that the first eigenvector (or principal component, PC) encompassed about 33% of the total phenotypic variance, which was significantly higher than expected under the null Wishart distribution (Fig. 4G). In other words, there was one dimension of shared evolutionary change that contained significant parallelism. We next examined if the different lake-stream systems evolve in parallel or anti-parallel directions by examining the loading of each PC, where a shared sign (positive or negative) is indicative of parallel evolution (*34*). The two *H. stappersii* and *P. philander* lake-stream systems had positive loadings, while the remaining seven lake-stream systems (all *A. burtoni* and *C. horei*) had a negative loading on the first PC. This indicates that all *A. burtoni* and *C. horei* systems evolve in parallel, but in an anti-parallel direction compared to *H. stappersii* and *P. philander.* Therefore, both parallel and anti-parallel evolution was detected in phenotypic divergence in haplochromine cichlid lake-stream systems. To infer which phenotypic characteristics were evolving in parallel among the different lake-stream systems, we examined which landmarks were contributing the most to the first PC by examining PC1 loading values. We found that the seven landmarks with the most extreme loading values (<-0.4 or >0.4) were related to mouth position (landmark y1), eye size (landmarks x4, y5) and slenderness of the body (body depth/standard length ratio and landmarks y7, y8, y9) (Fig. S5J, M).

We finally examined the genetic outlier data and found that the first PC encompassed about 30% of the total genetic variance, which was significantly higher than expected under the null Wishart distribution (Fig. 4J). Remarkably, the signs of PC1 loadings (positive or negative) were the same for each lake-stream system as for the phenotype data, namely positive for *H. stappersii* and *P. philander,* and negative for the remaining seven lake-stream systems. This further highlights parallel and anti-parallel evolution in lake-stream divergence. Among the eight PCs of the genetic outliers with the most extreme loading values (<-0.3 or >0.3), four PCs (PC5, PC7, PC10, PC11) were separating lake and stream populations (Fig. S5K,N). This indicates that half of the PCs contributing to genetic parallelism are involved in lake-stream divergence. Finally, when examining the genetic non-outlier data, we found that the first PC encompassed about 25% of the total genetic (non-outlier) variance, which was also significantly higher than expected under the null Wishart distribution (Fig. S5E). However, the interpretation of parallelism in the context of genetic non-outliers is less straightforward, as the seven genetic non-outlier PCs with the most extreme loading values (<-0.3 or >0.3) were not related to lake-stream divergence, but rather to species separation or to divergence within *H. stappersii* populations (Fig. S5L,O).

### Parallelism is higher when ancestral populations are more similar

We then examined whether the degree of similarity in the ancestral lake populations correlates with the extent of genetic and morphological parallelisms. We found a significant correlation in both datasets with a stronger effect in the genetic than in the morphological data (Mantel tests: *θ_G_outlier_* versus *F_ST_*: *P* = 0.0039; *θ_P_* versus *D_M_*: *P* = 0.045; *R^2^* = 0.46 and 0.17, respectively) (Fig. 4H, K), indicating that phenotypic and genetic parallelisms are higher when ancestral populations are phenotypically and genetically more similar. The genetic non-outliers (that is, the neutral markers) did not reveal such a correlation (Mantel test: *θ_G_* versus *F_ST_*: *P* = 0.48; Fig. S5F). Finally, we found that the proportion of standing genetic variation between ancestral populations was negatively correlated with *θ_P_* and *θ_G_outlier_* (Mantel tests: *P* = 0.0061 and 0.0117; *R^2^* = 0.32 and 0.43, respectively) (Fig. S5G, H) but not with *θ_G_* (Mantel test: *P* = 0.19; Fig. S5I), indicating that lake-stream population pairs sharing large amounts of standing genetic variation display more parallelism at the level of both phenotype and genotype.

### Body shape convergence in species inhabiting the same rivers

We finally assessed levels of convergence (and divergence) in phenotype and genotype across all lake-stream systems. We first examined how genomic differentiation scales with morphological differentiation across species. We thus contrasted the levels of absolute genomic (*d_XY_*) and morphological (*D_M_*) differentiation between all *A. burtoni* populations and those of the three other haplochromine species, resulting in 136 pairwise comparisons. We used *d_XY_* rather than *F_ST_* as *d_XY_* is an absolute measure of genomic differentiation that is better suited for between-species comparisons. As above, we classified these comparisons according to the environmental contrasts (lake-lake; lake-stream; stream-stream). The extent of body shape differentiation was only partially in agreement with the respective levels of genome-wide divergence (Fig. 4L). For example, the population pair from the Kalambo River involving the upstream population (*A. burtoni* Kalambo 2: *D_M_* = 5.6) was morphologically more distinct than the between-species comparisons in the Rusizi River (*A. burtoni* versus *H. stappersii*: *D_M_* = 4.1). In agreement with the measured levels of genomic differentiation (*d_XY_* = 3.6x10^-3^), morphological differentiation was high in the *P. philander* population pair (*D_M_* = 6.0).

We then calculated the among-lineage covariance matrices of mean trait values D_river_ and D_lake_ for each dataset (phenotype, genotype outlier, genotype non-outlier) to investigate levels of convergence/divergence in lake-stream differentiation. Following the definition of DeLisle & Bolnick (*34*), less variance in D_river_ compared to D_lake_ is indicative of convergent evolution, while more variance in D_river_ compared to D_lake_ is indicative of divergent evolution. We found divergence at the genomic level in both ‘outlier’ and ‘non-outlier’ datasets, since trace subtraction (tr(D_river_)-tr(D_lake_)) was positive in both cases. In contrast, we found convergence at the phenotypic level, since the result of trace subtraction was negative. In this latter case, two traits encompassed more than 99% of the total variance, namely centroid size (66%) and total length (33%). This might be explained by the fact that the different traits and landmarks have different units (e.g. variation in total length is in cm, while variation in landmarks is in mm), and therefore account differently for the total amount of variance (this inherent issue to the method has been identified by DeLisle & Bolnick (*34*), and references therein). Therefore, a more intuitive and biologically more meaningful way to assess convergence/divergence is to compare Euclidean distances for a specific trait. Thus, for each pair of lake-stream systems, we compared the Mahalanobis distance (*D_M_*) between the respective lake and the stream populations, where convergence is suggested when *D_M river_ -D_M lake_* < 0, with a more negative value indicative of higher convergence (*34*).

Within *A. burtoni*, the majority of pairwise comparisons indicated divergent evolution in body shape except for the pairs Chitili/Kalambo 1 and Chitili/Kalambo 2, with the highest level of convergence observed for the latter comparison (Table S9). The between-species comparisons revealed mostly divergent evolution, except for comparisons between the southern populations of *A. burtoni* and *C. horei*, which all indicated convergence. Remarkably, the highest level of convergence was found between *A. burtoni* and *C. horei* sampled in sympatry in the Lufubu River. Indeed, the lake populations are extremely differentiated (*D_M lake_* = 11.6), which was one of the most divergent comparisons in terms of morphological differentiation across the four species (Fig. 4L). In contrast, *A. burtoni* and *C. horei* from Lufubu River were morphologically very similar (*D_M river_* = 7.2) given their high level of genome-wide divergence (*d_XY_* = 8.5x10^-3^) (Fig. 4L) and similar levels of within-species lake-stream divergence (*A. burtoni*: *D_M_* = 4.8; *C. horei*: *D_M_* = 4.9) (Fig. 4M). Furthermore, body shape convergence was also observed in *A. burtoni* and *H. stappersii* populations from the Rusizi system. Indeed, while the Rusizi Lake populations are morphologically differentiated (*D_M lake_* = 5.4), the respective populations from Rusizi River are morphologically much more similar (*D_M river_* = 4.1), despite almost identical levels of genome-wide differentiation (*d_XY_* = 6.5x10^-3^ for both comparisons) (Fig. 4L, M, Table 2). Moreover, the level of between-species morphological differentiation in the same river (*D_M_* = 4.1) was smaller than the within *A. burtoni* differentiation in contrasting environments (*D_M_* = 4.6) (Fig. 4M, Table S3). Thus, we observed body shape convergence in species that co-occur in the same river systems and, hence, diversified along the very same environmental gradient. Finally, we can rule out gene flow between species as the reason for these similarities between species (Figs. S6, S7).

## Discussion

### Dynamics of genomic and morphological diversification in Astatotilapia burtoni

The first aim of this study was to investigate the dynamics of genomic and morphological diversification along the speciation continuum in the cichlid model species *Astatotilapia burtoni*. We sequenced 132 whole genomes and analysed the body shape of 289 *A. burtoni* individuals from six lake-stream populations pairs displaying increasing levels of genome-wide divergence to investigate how genomic and morphological differentiation accumulate along an ecological gradient and across geographic distances, in a setting where stream populations are derived from lake populations. We found that at early stages of differentiation, lake-stream population pairs displayed higher levels of genomic divergence than lake-lake population pairs for equivalent geographic distances. Furthermore, consistent with previous findings based on RAD-seq data (*25*), we found that stream populations have smaller effective population sizes than their respective lake population, possibly reflecting a founder effect and/or physical limitations of the riverine environment compared to the lake environment. Therefore, the combined effect of divergent selection and genetic drift likely accelerate genome-wide differentiation at early stages of divergence.

Furthermore, we assessed the dynamics of morphological differentiation by examining body shape differences among populations, as a slender body shape has been suggested to be adaptive in the stream environment (*24*). We found that morphology was the first axis of differentiation in both lake-stream and lake-lake comparisons compared to genomic differentiation and geographic distance. Then, differentiation trajectories diverged as genomic differentiation built up rapidly over short geographic distances in the presence of divergent selection and drift (i.e. in a lake-stream setting), while genomic differentiation built up more slowly in the presence of drift only (i.e. in a lake-lake setting). The relatively more rapid morphological differentiation in the early phases of diversification may be explained by the fact that variation in body shape is due to genetic and environmental factors (*24, 35*).

We established that divergent natural selection is a driver of early diversification in *A. burtoni* but whether or not it can ultimately lead to speciation remains an open question. We thus investigated how genomic and morphological differentiation build up across all *A. burtoni* populations, taking into account geographic distance. Our sample of population pairs spans a complete continuum of genomic divergence, from virtually panmictic populations to population pairs that resemble separate species (Fig. 2A). We found that morphological differentiation was gradual, while we observed a gap in genomic differentiation that could only partially be explained by geographic distance. While *F_ST_* values were not higher than 0.2 for geographic distances ranging from 1 to 40 km, there were two groups of *F_ST_* values for geographic distances ranging from 70 to 140 km; the ‘one-species’ (*F_ST_* <0.3) and the ‘two-species’ category (*F_ST_* >0.6). Our results in haplochromine cichlids are, thus, similar to what has been observed in *Timema* stick insects, in which comparable levels of genomic differentiation were found along an ecologically-driven speciation continuum (*F_ST_* < 0.3 in within-species population comparisons versus *F_ST_* > 0.7 in between-species comparisons) (*13*). Overall, our results in cichlids are in agreement with the ‘genome-wide congealing’ theory (GWC), proposing that in the presence of high levels of migration, divergent selection and linkage, there can be a ‘sudden’ transition from a state of well-intermixed populations to two reproductively isolated entities (i.e. a tipping point), due to a positive feedback loop between the levels of divergent selection and linkage disequilibrium (*3, 36*). We would like to note, however, that the lake-stream population pairs in the present study do not constitute an empirical example of the GWC theory, given that this theory has been developed based on a model of primary divergence with gene flow, while the majority of lake-stream population pairs are likely to have diverged under a scenario of secondary contact (i.e. including one or several periods of allopatric divergence). The observed demographic scenarios can be explained by the history of LT, which is characterized by lake level fluctuations (*37*) that may have isolated lake and stream populations when the lake level was low. Interestingly, our analyses revealed a zone of secondary contact in the Lufubu system, as shown by demographic modelling. This population pair displays intermediate levels of genomic differentiation (*F_ST_* = 0.47). The persistence over time of this population pair from the Lufubu River is unknown, as disentangling between transient stages (that is, a collapse in one species or a split in two species) and actual dynamic equilibria (that is, the presence of a hybrid zone) is unfeasible at present.

### Increasing levels of reproductive isolation along a continuum of genomic divergence

To assess to what extent genomic divergence scales with levels of reproductive isolation, we reviewed previous data that used the very same *A. burtoni* populations as in the present study and performed mate-choice experiments between the genetically most distinct *A. burtoni* populations from the North and the South of LT. Assortative mating was not detected in populations with low levels of genomic differentiation (*24, 27*), and we found that levels of reproductive isolation increased with genomic differentiation. Specifically, levels of reproductive isolation were already strong for parapatric and allopatric lake and stream populations with intermediate (*F_ST_* = 0.47) and high (*F_ST_* = 0.62) levels of genomic differentiation (*28*). Furthermore, mate choice experiments revealed signatures of partial assortative mating between the genetically divergent *A. burtoni* populations from the North and the South of LT (*F_ST_* = 0.69), but only in a setting including all cues (i.e. visual, olfactory and possibly acoustic). Consistent with this, it has previously been shown that olfactory cues are more important than visual cues in *A. burtoni* mate-choice (*38*). These results, together with their high levels of genomic divergence, suggest that the northern and southern *A. burtoni* populations behave like separate species, emphasizing that our sample of populations along a continuum of genomic divergence is representative of the entire speciation continuum. Taken together, previous data and our new experiment indicate that levels of reproductive isolation scale with levels of genome-wide divergence, which has been shown to be the case in many taxonomic groups (*39, 40*).

### A role of allopatry in the completion of speciation?

Considering the current distribution of the three most divergent lineages of *A. burtoni* (South of LT, North of LT, Lufubu River), it appears that the later stages of diversification could have been facilitated by periods of allopatry. Demographic modelling of the Lufubu lake-stream pair indicated that, while there is ongoing gene flow between the lake and the river populations, there was at least one period of allopatry between these populations that may have facilitated divergence. In addition, the most distant populations from the North and the South of LT are separated by more than 700 km, and phylogeographic work suggested that LT was colonized by the northern lineage from the Lukuga River (western part of LT), while the southern lineage colonized LT from the Lufubu River (South of LT) (*26*). Therefore, it is possible that these two lineages diverged before they colonized LT. However, the evolutionary history of *A. burtoni* is complex, and some hybridization events may have happened as indicated by shared mitochondrial lineages between populations from the North and the South of the lake, possibly linked to past lake-level fluctuations and/or long-distance migration events (*26*). For these two lineages, initial mechanisms of population divergence are unknown, but allopatry likely contributed towards the completion of speciation. This is consistent with the view that allopatry can promote and/or complete speciation (*2*).

### Rapid diversification in haplochromine cichlid fishes

We have shown that the *A. burtoni* population pairs with increasing levels of genomic divergence are representative of the speciation continuum, including populations with no assortative mating all the way to populations featuring partial assortative mating. We showed that strong levels of reproductive isolation are found at intermediate (*F_ST_* = 0.47) to high (*F_ST_* = 0.62; 0.69) levels of genome wide divergence, yet quite shallow levels of net nucleotide divergence (*D_a_*). Specifically, strong levels of reproductive isolation (here measured as extrinsic inviability) were measured for levels of *D_a_* ranging from 0.1% to 0.2%, and about 0.4% for premating isolation. However, isolation barriers appeared to have been porous at intermediate levels of genomic differentiation, since ongoing gene flow was measured in the Lufubu lake-stream pair at *F_ST_* = 0.47. Therefore, the “grey zone” of speciation in *A. burtoni* corresponds to *D_a_* ∼0.1% or *d_XY_* = 1.7x10^-3^. This suggests that, in haplochromine cichlids, reproductive isolation establishes at rather low levels of *D_a_* compared to what has previously been reported (*41*). More specifically, a study analyzing 63 populations or species pairs along a continuum of genomic divergence showed that the “grey zone” of speciation corresponded to *D_a_* levels ranging from 0.5% to 2% (*41*). However, we acknowledge that this study and our data are not entirely comparable due to different ways of measuring reproductive isolation (i.e. gene flow modelling (*41*) versus experimental measures in our study), and due to the fact that *D_a_* is a relative measure of genome differentiation depending on within-species genetic diversity. Therefore, the absolute measure of divergence *d_XY_* is better suited for between-species comparisons.

A recent study comparing patterns of genome divergence between species and population pairs of *Pungitius* sticklebacks showed that high genomic *F_ST_* values corresponded to very low estimated measures of gene flow (*15*). Furthermore, allopatric population pairs of *Pungitius sinensis* (*d_XY_*: 6.2-9.8x10^-3^) have similar or higher levels of absolute divergence compared to *A. burtoni* ‘North’ vs. *H. stappersii* (*d_XY_*: 6.4-6.5x10^-3^) or compared to *A. burtoni* ‘North’ vs. *A. burtoni* ‘South’ (*d_XY_*: 3.0-3.2x10^-3^). These low levels of between-species absolute divergence appear to be general in cichlids, as shown in 73 species representative of the recent Lake Malawi radiation where the average absolute divergence was 2x10^-3^ (*d_XY_* range: 1.0-2.4x10^-3^) (*42*). A similar range of *d_XY_* values has been reported in two ‘young’ (‘Python’ *d_XY_*: 2.05x10^-3^) and ‘old’ (‘Makobe’ *d_XY_*: 2.17x10^-3^) *Pundamilia* species pairs from Lake Victoria (*43*). Taken together, these results are consistent with rapid speciation and explosive diversification characteristic of cichlid fishes (*20*).

### Low sharing of regions of differentiation among lake-stream systems

The third aim of this study was to examine and quantify levels of (non-)parallelism among six *A. burtoni* lake-stream systems and three lake-stream systems from additional haplochromine species. To this end, we identified regions of differentiation that we defined as the overlap of the top 5% of *F_ST_*, *d_XY_* and π-difference values. We found between 2 and 101 such outlier regions ranging from 10 to 70 kb. The number of outlier regions reported here is relatively small compared to other studies in cichlids (*43, 44*), which, however, can be explained by the more stringent definition of such regions in our study (the intersection between three metrics). Furthermore, we found little overlap among differentiation regions from different lake-stream systems, as only 19 outlier regions were shared among two lake-stream systems, and not a single such region was shared by more than two systems. Interestingly, however, Bayesian analyses of selection revealed that *H. stappersii* and the southern *A. burtoni* populations shared a common set of overrepresented outlier loci involved in sensory perception (sound and light). Taken together, these results highlight that, although a large majority of outliers are not shared, some functions are important for riverine adaptation and may repeatedly be the target of natural selection. Consistent with this, a previous study (*43*) identified common highly differentiated genomic regions between a young and an old cichlid species pairs diverging along a depth gradient in Lake Victoria, and found that two thirds of the differentiation regions were private to each species pair. This highlights that adaptive divergence often encompasses both parallel and system-specific (non-parallel) components.

The overall low levels of gene and function sharing between lake-stream systems reported here may be due to cryptic environmental heterogeneity in the stream environment. The streams from which the populations of this study were collected are not only different in size but also encompasses diverse ecological niches that may require specific mechanisms of adaptation (e.g. (*33*)). A non-mutually exclusive explanation for the lack of sharing of regions of differentiation is that these regions may not be due to divergent selection but due to genetic drift in allopatry and background selection (*45*), as the majority of population pairs did not follow a scenario of primary divergence with gene flow. However, the background selection scenario is realistic mainly for population pairs with elevated levels of genome-wide differentiation (*46*). Finally, adaptive phenotypic plasticity (*27, 47*) as well as epigenetic factors (*48, 49*) might play complementary roles in adaptive divergence. Yet, their investigation was beyond the scope of the present study.

### Parallelism revealed by multivariate analyses

To better quantify the level of (non-)parallelism among the nine lake-stream systems, we performed Phenotypic Change Vector Analyses (PCVA) (*19, 32*). Pairwise examination of *_θ_P* and *θ_G_* _outlier_ revealed that diversification was not particularly parallel among lake-stream systems, as the majority of *_θ_P*- and *θ_G_* _outlier_-values were close to orthogonal. These values are similar to what has been previously reported in lake-stream population pairs of threespine sticklebacks (*33*). Interestingly, *θ_G_* _outlier_-values correlated with *_θ_P*-values, but *θ_G_* _non-outlier_-values did not. This strongly suggests that adaptive phenotypic divergence has a genetic basis in the species under investigation, which is similar to what has been reported for lake-stream population pairs of threespine stickleback fish (*33*).

It has recently been suggested that signatures of parallelism can be overlooked when relying on pairwise comparisons only (*34*). Therefore, we conducted the recently proposed multivariate analysis of the C matrix (*m* lake-stream systems x *n* traits), which is essentially a PCA of (phenotypic or genomic) vectors of differentiation to uncover major axes of evolutionary change (*34*). We found that for both phenotypic and genomic outlier datasets, there was one significant axis of parallel evolutionary change, in which seven out of nine lake-stream systems evolved in parallel. Remarkably, these seven lake-stream systems were the same in the phenotypic and genomic outlier datasets. In other words, all *A. burtoni* and *C. horei* lake-stream systems have a common major axis of phenotypic and genomic parallel evolution, while *H. stappersii* and *P. philander* were evolving in an anti-parallel manner with regards to the seven other lake-stream systems. Interestingly, the traits underlying parallel phenotypic evolution were related to mouth position, eye size and slenderness of the fish body, which have previously been suggested to contribute to adaptive divergence in *A. burtoni* (*24*). We show here that these phenotypic traits not only evolved in parallel in all *A. burtoni* lake-stream systems, including the Rusizi system from the North of LT, but also in *C. horei* from the Lufubu system. These traits are relevant for a stream-adapted life style, as a lower mouth position likely evolved in response to riverine trophic conditions and a slenderer body has likely evolved in adaptation to fast water flow conditions (*24*). Furthermore, eight PCs contributing most to parallel genotypic evolution were related to lake-stream divergence (4 PCs), species differences (3 PCs), and geographic divergence within *A. burtoni* (1 PCs). Therefore, the genomic outliers mostly encompass candidate loci for lake-stream genomic adaptive divergence. Surprisingly, the analysis of the genomic non-outliers also revealed a significant axis of parallel evolution. However, the genomic traits underlying parallelism were not related to lake-stream divergence, but rather summarized between species (1 PC), geographic (3 PCs) or within-population (3 PCs) divergence. Thus, the biological interpretation of this major axis of parallel evolution is less straightforward. To date, these multivariate analyses have been performed here and on a threespine stickleback dataset of 16 lake-stream population pairs (*34*), therefore it would be interesting to apply these analyses to a broader range of model systems to uncover additional major axes of parallel evolutionary changes.

### The distance between ancestral populations influences the level of parallelism

It has previously been suggested that the probability of parallelism at the molecular level decreases with time since divergence (*50*). Furthermore, it has been suggested that the extent of parallelism should be higher when ancestral populations were closely related (*17*). We thus examined if there was a correlation between the distances between ancestral (i.e. lake) populations and the level of parallelism (i.e. *θ*). We found that for both phenotypic and genetic outliers, there was a significant positive correlation between these metrics (*_θ_P* and *D_M_*; *θ_G_* _outlier_ and *F_ST_*), confirming that the morphological and genetic distances between ancestral populations influence the level of parallelism.

Standing genetic variation is an essential component of replicated adaptive evolution (*51*). Thus, we tested if the amount of standing genetic variation between ancestral populations was correlated to levels of parallelism in the respective lake-stream systems. In line with our predictions, we found that the levels of standing genetic variation and parallelism were negatively correlated. In other words, lake-stream population pairs sharing larger amounts of standing genetic variation display more parallelism at the level of both the phenotype and the genotype. Parallelism is, thus, likely constrained by the amount of standing genetic variation upon which natural selection can act, as the effect of *de novo* beneficial mutations on parallel evolution is much less likely to play an important role in adaptive divergence in recently diverged population pairs. Previous theoretical work has further shown that even small differences in the directionality of selection can greatly reduce genetic parallelism, especially in the case of complex organisms with many traits (*52*). This suggests that, besides time since divergence, also cryptic habitat heterogeneity (leading to small differences in the directionality of selection) can decrease the likelihood of parallelism. In support of this, a comparison of regional (within Vancouver Island) versus global (North America versus Europe) lake-stream population pairs of sticklebacks showed that parallelism decreases at increased spatial scales (*53*).

### Body shape convergence in species inhabiting the same rivers

Finally, we investigated levels of convergence or divergence among the nine lake-stream population pairs. While we found divergence at the genomic levels (for both outlier and non-outlier datasets), we found convergence at the morphological level by examining lake and river among-lineage covariance matrices of trait mean values. As body shape contributed significantly to the overall variance, we further focused on this trait for pairwise comparisons of morphological convergence/divergence. Remarkably, we found body shape convergence in both species pairs from the same river systems, namely *A. burtoni* and *H. stappersii* in the Rusizi River, and *A. burtoni* and *C. horei* in the Lufubu River (also note that the sympatric population-pairs show rather similar *F_ST_* values across species; Fig. 3). This highlights that local ecological selection constraints body shape evolution in haplochromine cichlids. This has previously been shown in Midas (*Amphilophus* sp.) cichlid fishes, where the body shapes of two syntopic species were more similar to each other than the body shape average of the first species from different localities (*54*).

That we did not find convergence at the genomic level despite convergence at the morphological level can arise from several non-mutually exclusive factors. First, the outlier detection method based on lake-stream differences might not be the most appropriate to uncover the genomic basis of morphological evolution. A quantitative traits loci approach might be more appropriate, as has previously been applied to uncover genomic regions underlying body shape differences along a benthic-limnetic axis of differentiation in Midas cichlids (*55, 56*). Second, while body shape convergence may be due to adaptive genomic divergence, adaptive phenotypic plasticity also plays a role in body shape evolution. Indeed, it has previously been shown that body shape is partially plastic in *A. burtoni* (*24*), and that adaptive phenotypic plasticity plays an important role in *A. burtoni* lake-stream divergence (*27, 47*). Further studies should focus on quantifying the respective influence of genomic divergence *versus* phenotypic plasticity in adaptive divergence.

## Conclusion

Our results reveal that diversification in the East African cichlid *A. burtoni* likely occurred under the influence of both divergent selection and geographic isolation, highlighting the combined roles of ecological and non-ecological processes in speciation. Furthermore, we found that morphological diversification was gradual along the speciation continuum, likely due to the fact that morphology is the result of the interaction of genetic and environmental factors. Contrastingly, we found a gap in genomic differentiation, providing support for the hypothesis that there is a tipping point in genomic differentiation during the speciation process. Furthermore, our study provides an empirical example of fast diversification inherent to cichlids, exemplified by a grey zone of speciation at shallow levels of genomic divergence. Additionally, the quantification of parallelism in nine lake-stream population pairs from four cichlid species revealed that while pairwise comparisons failed to identify strong signatures of parallelism, multivariate analyses allowed to uncover major axes of shared evolutionary changes along the lake-stream ecological gradient. To conclude, our study highlights that diversification is a complex product of differentiation trajectories through multivariate space and time.

## Materials and Methods

### Study Design

### Research objectives

In this study we investigated the dynamics of morphological and genomic diversification in *Astatotilapia burtoni* populations that have diverged along a lake-stream environmental gradient and across geography. We further aimed to investigate the extent and predictability of (non-)parallelism and convergence in nine lake-stream population pairs from four East African cichlid species.

### Research subjects or units of investigation

Four haplochromine cichlid species: *Astatotilapia burtoni, Haplochromis stappersii, Ctenochromis horei* and *Pseudocrenilabrus philander*.

### Experimental design

Individuals of *Astatotilapia burtoni* (N = 132), *Haplochromis stappersii* (N = 24), *Ctenochromis horei* (N = 24) and *Pseudocrenilabrus philander* (N = 24) were collected in Zambia and Burundi between January and November 2015 (Tables S1, S2). All fish were sampled with a ∼1:1 sex ratio and were adult specimens except for 3 *P. philander* juveniles from the Mbulu River. Fish were sampled in six different tributaries to LT, whereby each system comprises a riverine population (N = 10-12) and a lake population (N = 12-14) and was named after the river, except for the Lake Chila system that was sampled outside of the LT basin. *H. stappersii* were sampled at the Rusizi River, in the North of LT, along with sympatric *A. burtoni* populations (Fig. 1; Table S1). All other populations were sampled in the South of LT. *A. burtoni* and *C. horei* were sampled at the Lufubu River; *A. burtoni* was further sampled in the Lunzua, Chitili and Kalambo rivers (Fig. 1; Table S1). As two river populations were sampled in the Kalambo River, two lake-stream population comparisons were used for this river, namely Kalambo1 (comparison Kalambo lake versus Kalambo1) and Kalambo2 (comparison Kalambo lake versus Kalambo2). Finally, *P. philander* were sampled in small Lake Chila and in Mbulu creek (Fig. 1; Table S1). All fish were caught with fishing rods or minnow traps and anaesthetised using clove oil. Photographs of the leſt lateral side were taken using a Nikon D5000 digital camera, under standardised lighting conditions, and with a ruler for scale. To aid in digital landmark placement, three metal clips were used to spread the fins at the anterior insertions of the dorsal and anal fin, and at the insertion of the pectoral fin (Fig. S2A). To increase the sample size for morphological analyses, additional individuals were sampled and photographed at the same locations and time points as the individuals whose genomes were sequenced (Table S2). Standard length, total length, and weight were measured. A piece of fin clip was preserved in 99% ethanol for DNA extraction. Whole specimens were preserved in 70% ethanol.

### Sample size

For the genomic analyses, we planned to sample 12 individuals per population (6 males and 6 females) as 24 alleles per population are sufficient to obtain accurate allele frequency estimates. We sampled only 10 *Astatotilapia burtoni* specimens from the Rusizi River because we did not succeed to catch additional specimens after numerous attempts. 14 *A. burtoni* specimens from Rusizi lake were sampled to obtain a total of 24 *A. burtoni* specimens from the Rusizi lake-stream system. For the morphometric analyses, we intended to photograph at least 17 specimens per population, which is equal to the number of landmarks used. Fewer specimens were photographed in three populations (*A. burtoni* Chitili lake, N=13; *A. burtoni* Rusizi River, N=10; *P. philander* Mbulu Creek, N=9) because we did not succeed to catch additional specimens after numerous attempts and because 3 of the individuals caught in Mbulu Creek were juveniles and thus excluded from the morphometric analyses.

### Data inclusion/exclusion criteria and outliers

For the genomic analyses, all individuals were included in the analyses except one hybrid specimen between *A. burtoni* and another species that was detected only after whole genome sequencing. For the morphometric analyses, juveniles were excluded on the basis that their morphology is different from the adults’ morphology.

### Replicates

The first mate-choice experiment (including only visual cues) was replicated 44 times; each replicate included one trio of fish (1 female, 2 males from different populations). The second mate-choice experiment (including direct contact) was replicated 8 times; each replicate included 8 fish (2 males and 6 females). Additional replicates could not be performed due to fish number limitations in the laboratory.

### DNA extraction, sequencing, data processing

DNA was extracted from fin clips using the EZNA Tissue DNA Kit (Omega Bio-Tek) following the manufacturer’s instructions. Individual genomic libraries were prepared using TruSeq DNA PCR-free Low Sample Kit (Illumina) and subsequently sequenced (150 bp paired-end) on an Illumina HiSeq3000 sequencer at the Genomics Facility Basel (GFB) operated jointly by the ETH Zurich Department of Biosystems Science and Engineering (D-BSSE) and the University of Basel.

For each library, the quality of raw reads was visually inspected using FastQC (v0.11.3) and Illumina adapters were trimmed using Trimmomatic (*57*) (v0.36). Filtered reads of each individual were aligned separately against the *Metriaclima zebra* reference genome (assembly M_zebra_UMD1). We chose this reference genome rather than the *Astatotilapia burtoni* reference genome (*23*) to avoid any reference bias when comparing *A. burtoni* with the other species. We also chose *M. zebra* rather than *Oreochromis niloticus* as reference genome to maximise the number of reads mapped, as *M. zebra* is phylogenetically closer to *A. burtoni, C. horei, H. stappersii*, and *P. philander* than *O. niloticus*.

The *M. zebra* reference genome was indexed using BWA (*58*) (v.0.7.12) and alignments were performed using BWA-mem with default parameters. Obtained alignments in SAM format were coordinate-sorted and indexed using SAMtools (*59*) (v.1.3.1). The average coverage per individual ranged from 9.8× to 24.5× (Table S1). We performed an indel realignment using RealignerTargetCreator and IndelRealigner of the Genome Analysis Tool Kit (GATK) (*60*) (v3.4.0). Variants were called using the GATK functions HaplotypeCaller (per individual and per scaffold), GenotypeGVCFs (per scaffold), and CatVariants (to merge all VCF files). The VCF file corresponding to the mitochondrial genome (scaffold CM003499.1) was then isolated from the VCF file corresponding to the nuclear genome (that is, all other scaffolds). The latter was annotated with the features ExcessHet (that is, the Phred-scaled p-value for an exact test of excess heterozygosity) and FisherStrand (that is, the Strand bias estimated using Fisher’s exact test) using the GATK function VariantAnnotator. To filter the VCF file, empirical distributions of depth (DP) and quality (QUAL) were examined. The VCF file was filtered using the GATK function VariantFiltration with the following values (variants meeting the criteria were excluded): DP<2000; DP>4000; QUAL<600; FisherStrand>20; ExcessHet>20.

In addition, variants were called using SAMtools mpileup (per scaffold) with the following options: - C50 -pm2 -F0.2 -q 10 -Q 15. Files per scaffold were then converted to VCF format, concatenated (except the mitochondrial genome) and indexed using BCFtools (v.1.3.1). The VCF file was annotated for ExcessHet and FisherStrand, and the distribution of depth and quality were visually assessed as described above. The VCF file was filtered using the GATK function VariantFiltration with the following values: DP<1500; DP>4000; QUAL<210; FisherStrand>20; ExcessHet>20. Filtered GATK and SAMtools datasets were then combined using bcftools isec. The final VCF file contained variants present in both datasets. Genotypes were then imputed and phased per scaffold using beagle (v.4.0). In total, the final VCF file contained 26,704,097 variants. Chomonomer (v.1.05; http://catchenlab.life.illinois.edu/chromonomer/) was used to place the 3,555 *M. zebra* scaffolds in 22 linkage groups using two linkage maps (*61*). For BayPass selection analyses and allele frequency calculations (see below), we excluded indels and non-biallelic sites, resulting in a VCF file containing 20,343,366 SNPs.

### Genetic structure and phylogenetic relationships

Population genetic structure was examined using principal components analyses (PCA) implemented in the smartPCA module of Eigensoft (v.6.1.1). To reconstruct a whole-genome nuclear phylogeny, a sequence corresponding to the first haplotype of each scaffold was extracted using bcftools consensus --haplotype 1 of BCFtools v1.5 (https://github.com/samtools/bcftools). Individual whole genome sequences were then concatenated and a maximum-likelihood (ML) analysis was performed in RAxML (*62*) (v.8.2.11) using the GTRGAMMA sequence evolution model and 20 fast bootstrap replicates. The option –*f a* was used to report the best-scoring ML tree with branch lengths. KBC4, a putative *A. burtoni* individual sampled in the Rusizi River, did not cluster with other *A. burtoni* individuals in the phylogeny (labelled “hybrid” in Fig. S1A). This specimen results most likely from a hybridisation event with *Astatoreochromis alluaudi,* as its mitochondrial genome is closely related to *A. alluaudi* (data not shown). Therefore, this individual was excluded from further analyses. In order to test for introgression or retention of ancestral polymorphism between sympatric species (that is, *A. burtoni* and *H. stappersii* in the Rusizi system, and *A. burtoni* and *C. horei* in the Lufubu system), a topology weighting analysis reconstructing fixed-length 5-kb-windows phylogenies was performed using *Twisst* (topology weighting by iterative sampling of subtrees (*63*)).

### Demographic modelling

For each of the nine lake-stream population pairs, demographic simulations based on the joint site frequency spectrum (SFS) were performed in order to estimate the most likely model of population divergence as well as the best values of demographic parameters (effective population sizes, divergence times, migration rates). Simulated SFS were obtained using diffusion approximation implemented in ∂a∂I (*64*) (v.1.7.0). A modified version of the program including additional predefined models and the calculation of the Akaike Information Criterion (AIC) for model selection was used for the simulations (*65*). Eight demographic models of population divergence were tested (Fig. S3A): Bottle-Growth (BG), Strict Isolation (SI), Isolation with Migration (IM), Ancient Migration (AM), Secondary Contact (SC), as well as versions of these models including two categories of migration rates (IM2M; AM2M; SC2M). These two categories of migration rates can separate, for example, selected versus neutral loci. For the population pairs with a genome-wide *F_ST_* > 0.47 (*A. burtoni* Lufubu, *C. horei* Lufubu, and *P. philander*), the model BG was not tested as it was obvious that the populations are separated. Each model was fitted to the observed joint SFS using three successive optimisation steps: “hot” simulated annealing, “cold” simulated annealing, and BFGS (*65*). For each lake-stream population pair, 20 replicate analyses comparing seven or eight models were run, using different parameter starting values to ensure convergence. After these 20 runs, the model displaying the lowest AIC and the least variance among the replicates was chosen as the best model. For parameter estimation, additional runs were performed so that the total number of runs was 20 for the best model. To calculate the divergence times in years, a generation time of one year was used. As the scaled mutation rate parameter *θ* is estimated, we used the relation *θ* = 4×Ne×µ×L to infer the ancestral effective population size (*Ne*). The mutation rate (*µ*) (3.51x10^-9^ mutation per generation per year) of Lake Malawi cichlids (*42*) and the length of the genome assembly (L) of *M. zebra* (UMD1: 859,842,111 bp) were used.

### Regions of differentiation

For each lake-stream system, genome-wide *F_ST_* (Hudson’s estimator of *F_ST_*), | π_lake_ –π_stream_ | (absolute value of the difference in nucleotide diversity between the lake and the river populations), and *d_XY_* (absolute divergence) were calculated on 10 kb non-overlapping sliding windows using evo (https://github.com/millanek/evo). We defined as window of differentiation each window that is contained in the overlap of the top five percent values of these three metrics. Adjacent differentiation windows and windows separated by 10 kb were then merged in differentiation regions. To test if the differentiation regions of each system are affected by chromosome centre-biased differentiation (CCBD (*30*)), each region was placed either in the “centre” or in the “periphery” categories. These categories were defined by splitting each chromosome into four parts of equal length, where the “centre” category encompasses the two central parts of the chromosome and the “periphery” category encompasses the two external parts of the chromosome.

### Bayesian selection and association analyses

To detect signatures of selection at the SNP level, we used the Bayesian method BayPass (*31*). The core model performs a genome scan for differentiation by estimating a population covariance matrix of allele frequencies. It allows determining outlier SNPs based on the top 1% of simulated *XtX* values, where *XtX* is a differentiation measure corresponding to a SNP-specific *F_ST_* corrected for the scaled covariance of population allele frequencies (*31*). As the northern and southern populations of *A. burtoni* are highly divergent (see (*26*) and Fig. S1), only the southern *A. burtoni* populations were analysed jointly. We thus compared outlier sets for the southern populations of *A. burtoni*, the northern populations of *A. burtoni,* and *H. stappersii*. For the southern *A. burtoni* populations, an additional association analysis using one categorical covariate (lake population versus stream population) was performed using the auxiliary variable covariate model (AUX). Five replicate runs were performed using different starting seed values and default search parameters, except for the number of pilot runs (25). The final correlated SNP set contained the overlap of SNPs for which the Bayes Factor (BF) was higher than 10 and which were in the top five percent of δ values (the posterior probability of association of the SNP with the covariable) in the five replicate runs.

### GO annotation and enrichment analyses

To infer if candidate genes in differentiation regions were enriched for a particular function, all genes included in differentiation regions of each lake-stream system were extracted. In addition, genes including overlapping SNPs between the southern *A. burtoni,* the northern *A. burtoni*, and *H. stappersii* core outliers were reported, as well as genes including overlapping SNPs between *A. burtoni* core outliers and SNPs significantly correlated with lake versus stream environment and morphology. All candidate genes were blasted (blastx) against the NR database (version 12.10.2017) using BLAST+ v.2.6.0 and the first 50 hits were reported. To obtain a reference gene set, all *M. zebra* genes were blasted against NR in the same way. Gene Ontology and InterProScan annotations were retrieved from Blast2GO PRO (v.4.1.9). Enrichment analyses were performed using Fisher’s exact test for each differentiated gene set (one set per system) against the reference gene set (significance level: 0.001). For the genes located in the overlap of differentiation regions among systems, an additional step was performed by manually retrieving the annotations from *Homo sapiens* dataset in Uniprot (accessed online 20.11.2017).

### Diversification dynamics and genomic divergence in similar versus contrasted environments

To infer the dynamics of genomic differentiation along the lake-stream axis, genome-wide pairwise *F_ST_* and the net nucleotide difference *D_a_* (proxy of time since differentiation; *d_XY_* –(π_1_+π_2_ / 2)) were calculated for all possible population pair combinations, resulting in 136 within and between systems comparisons. A logarithmic regression was fitted to the data using the lm function (lm(*F_ST_*∼ln(*D_a_*) implemented in R (*66*) (v.3.4.2). To estimate the influence of divergent selection at early stages of genomic differentiation in sympatry/parapatry, population pairwise *F_ST_* of the southern populations of *A. burtoni* were used (‘lake-lake’: 6 comparisons; ‘lake-stream’: 5 comparisons), as well as the pairwise Mahalanobis distances (*D_M_*; see below). The northern *A. burtoni* populations were not used due to the high levels of genomic divergence compared to the southern populations. For each comparison, the geographic coastline distance between populations was measured using Google Earth (https://www.google.com/intl/en/earth/). Then, a linear model was fitted for each category (‘lake-lake’ and ‘lake-stream’) and the adjusted coefficient of determination *R^2^* was reported in R.

### Morphometric analyses

Geometric morphometrics was used to compare adult body shape between populations. Three juvenile individuals of *P. philander* from Mbulu creek whose genomes had been sequenced were excluded from the morphological analyses. In total, the photographs of 468 individuals (Table S2) were used for geometric morphometric analyses (289 *A. burtoni*, 81 *H. stappersii*, 67 *C. horei*, and 31 *P. philander)*. Using TPSDIG2 (*67*) (v.2.26) we placed 17 homologous landmarks (Extended Data Fig. 2a) on the lateral image of each fish. The tps file with x and y coordinates was used as an input for the program MORPHOJ (*68*) (v.1.06d) and superimposed with a Procrustes generalized least squares fit (GLSF) algorithm to remove all non-shape variation (i.e. size, position and orientation). Additionally, the data were corrected for allometric size effects using the residuals of the regression of shape on centroid size for further analyses. Canonical variate analysis (CVA) (*69*) was used to assess shape variation among *A. burtoni* populations (Fig. S2B,D) and among all populations of the four species (Fig. S2C,E). The mean shape distances of CVA were obtained using permutation tests (10,000 permutations). Mahalanobis distances (*D_M_*) among groups from the CVA were calculated and plotted against genetic distances (*F_ST_* for within *A. burtoni* comparisons; *d_XY_* for between species comparisons).

### Vectors of phenotypic and genomic divergence

We followed the method first developed by Adams and Collyer (*32*) and described in detail in Stuart et al. (*33*) and Bolnick et al. (*19*) to calculate multivariate vectors of phenotypic and genomic divergence. For vectors of morphological divergence, 37 traits and landmarks were used: centroid size (17 landmarks), standard length (landmarks 1-14), body depth (landmarks 8-12) corrected by standard length (ratio BD/SL), and the x and y coordinates of each of the 17 landmarks. We then calculated two types of vectors of genomic divergence. First, ‘outlier’ vectors were calculated using the first 78 principal components (88% variance; same as for non-outliers) of a genomic PCA based on outliers SNPs present in at least one set of BayPass outliers (*H. stappersii* core model; *A. burtoni* North core model; *A. burtoni* South core model; *A. burtoni* South auxiliary model). Then, ‘neutral’ vectors were calculated using the first 78 principal components of a genomic PCA based on all remaining SNPs (that is, non-outliers) (88% variance; all PC summarising between-population variation). For each lake-stream pair and separately for phenotypic and genomic outlier and non-outlier data, we calculated vector length (*L*), the difference in length between two vectors (*ΔL*) and the angle in degrees between two vectors (*θ*). For each morphological trait (respectively each genomic PC), we ran *t-*tests and used the *t-*statistic as an estimate of lake-stream divergence for each trait. Thus, vectors of phenotypic divergence were represented as the matrix C*_P_* of 37 *t-*statistics for each trait/landmark (columns) × 9 lake-stream pairs (rows), and vectors of genomic divergence were represented as a matrix C*_G_* of 78 *t-*statistics for each PC × 9 lake-stream pairs. We calculated 9 *L_P_* (phenotype), 9 *L_G_* (genotype), 9 *L_G_outlier_* values, all lake-stream pairwise comparisons *ΔL_P_*, *θ_P_*, *ΔL_G_*, *θ_G_*, *ΔL_G_outlier_* and *θ_G_outlier_*. We used Mantel tests (mantel.test function in the ape R package v.5.3; 9,999 permutations) to test the correlation between: *θ_P_* and *θ_G_ / θ_G_outlier_*; *ΔL_P_* and *ΔL_G_ / ΔL_G_outlier_* and linear regression models (lm function in R) to test the correlation between *L_P_* and *L_G_ / L_G_outlier_*.

### Multivariate analyses of (non-)parallelism and convergence

In addition to comparing all lake-stream divergence vectors in a pairwise manner, we performed the recently described eigen analyses of vector correlation matrices (C) (*34*). These multivariate analyses allow to quantify the extent of parallelism/anti-parallelism by assessing the percentage of variance explained by the leading eigenvectors of C. These analyses also reveal how many dimensions of shared evolutionary change exist, by inferring how many eigenvalues are significant. For each dataset (phenotype, genotype ‘outlier’ and genotype ‘non-outlier’), we calculated the eigen decomposition of the respective matrix of vector correlations C described above (C = QVQ^-1^) to extract eigenvectors (Q) and eigenvalues (V) of each dataset. To construct a null expectation of evolutionary parallelism, we sampled the *m*-dimensional Wishart distribution with *n* degrees of freedom as suggested by De Lisle & Bolnick (*34*), where *m* is the number of lake-stream pairs and *n* is the number of traits/landmarks. To infer which traits/landmarks/PCs contribute the most to the leading eigenvector of C, we investigated the value of trait loadings in the matrix A, where A = X^T^Q^T -1^.

Furthermore, we investigated levels of convergence/divergence in multivariate trait space by comparing the among-lineage covariance matrices of trait mean values (D) for each environment (*34*). Specifically, for each one of the three datasets (phenotype, genotype ‘outlier’ and genotype ‘non-outlier’), we calculated D_river_ and D_lake_ and their respective trace which encompasses the total among-lineage variance per environment. To assess levels of convergence/divergence in lake-stream divergence, we then compared both trace values (tr(D_river_)- tr(D_lake_)), where a negative value indicates convergence (less among-lineage variance in the river environment) whereas a positive value indicates divergence (more among-lineage variance in the river environment). Finally, it has also been proposed to investigate convergence/divergence of a specific trait by estimating the change in Euclidean distance between lineage pairs from different environments (here river vs. lake) (*19, 34*). We therefore calculated, for each pair of lake/stream system, the difference in Mahalanobis distance between the respective stream population pair vs. lake population pair. For instance, the convergence/divergence in body shape between *A. burtoni* and *C. horei* from the Lufubu system was calculated as follow: (*D_M A.burtoni_* _Lufubu river – *C.horei* Lufubu river_) - (*D_M A.burtoni_* _Lufubu lake – *C.horei* Lufubu lake_); where a negative value indicates convergence and a positive value indicates divergence in body shape.

### Predictability of (non-)parallelism

We then investigated the influence of similarity of ancestral (i.e. lake) populations on parallel evolution. We used Mahalanobis distances and *F_ST_* between lake populations as proxies of morphological and genetic similarities, respectively. We performed Mantel tests between lake-lake Mahalanobis distances and *θ_P_*, and between lake-lake *F_ST_* and *θ_G_ / θ_G_outlier_* to infer if similarity of ancestral populations between systems was correlated to the direction of (non-)parallelism. Finally, we assessed if the proportion of standing genetic variation is correlated with the extent of morphological or genetic parallelism. For each of the 136 population pairwise comparison, we extracted allele frequencies of biallelic SNPs using VCFtools --freq command. We then categorised SNPs in four different groups: identical sites (fixed in both populations for the same allele); differentially fixed sites (fixed in both populations for different alleles); fixed and variable sites (fixed in one population and variable in the second population) and standing genetic variation (SGV; variable in both populations). We performed Mantel tests between the proportion of SGV SNPs between lake populations and *θ_P_*, and between the proportion of SGV SNPs and *θ_G_ / θ_G_outlier_*.

### Mate-choice experiments

We used two mate-choice experiments to test for reproductive isolation between the two geographically most distant and genetically most divergent *A. burtoni* populations, Rusizi lake and Kalambo lake. Detailed methods and results of the experiments are provided in Appendix 1. Briefly, in the first experiment, a two-way female choice set up was used to test whether females preferred males of their own population over others when only visual cues are available (Fig. S4A). We placed a gravid female of either population (*N* = 44) in a central tank and allowed visual contact and interaction with two sized-matched males from Rusizi lake and Kalambo lake presented in two outer tanks. Within a period of up to 12 days, we assessed if the female had laid the eggs in front of the Rusizi lake male, the Kalambo lake male, in front of both, or in the central section. The position of the laid eggs was used as a measure for female preference (conspecific choice coded as 1; heterospecific and no choice coded as 0). The binomial data were then analysed with a generalized linear mixed model, which tested if the probability of the females spawning with the conspecific male was significantly different from 0.5.

In the second experiment, female spawning decisions were determined in a multi-sensory setting with free contact between females and males. A single tank was subdivided into three equally sized compartments by plastic grids (Fig. S4B). The middle compartment offered a resting and hiding place for the females whereas the two outer compartments served as male territories. The grid size was chosen to allow the smaller females to migrate between the three compartments, and to prevent direct contact between the larger males to exclude male-male competition. We conducted eight trials, each time using two males (one of each population) and six females from both populations (*N*_males_ = 16; *N*_females_ = 46). Mouthbrooding females were caught and ten larvae from each were collected for paternity analyses based on five microsatellite markers. All adult males and females were also genotyped for these five markers. The percentage of fertilised offspring by con- or heterospecific males in each replicate was then used to infer if the females spawned more frequently with the conspecific males. These experiments were performed under the cantonal veterinary permit nr. 2356.

### Statistical Analysis

Statistical parameters including the exact value of *N* are reported in the methods and figure legends. All statistical analyses were performed using R v.3.4.2.

## Acknowledgments

**General**: We would like to thank Fabrizia Ronco, Adrian Indermaur, and Gaëlle Pauquet for their help during fieldwork, the crews of the Kalambo Lodge and the Ndole Bay Lodge for their logistic support in Zambia, and the Lake Tanganyika Research Unit, Department of Fisheries, Republic of Zambia, and the University of Burundi, the Ministère de l’Eau, de l’Environnement, de l’Amenagement du Territoire et de l’Urbanisme, Republic of Burundi, for research permits. We thank Athimed El Taher and Milan Malinsky for their help with data analyses, Matthew Conte for sharing the chromonomer results, Astrid Böhne and Michael Matschiner for helpful comments on the manuscript, and Julie Johnson for fish illustrations. Finally, we are grateful to the support team of sciCORE (center for scientific computing, University of Basel, http://scicore.unibas.ch/) for providing access to computational resources, especially Pablo Escobar Lopez.

**Funding**: This study was supported by grants from the Swiss Zoological Society (SZS) to JR and the Swiss National Science Foundation (SNSF, grant 31003A_156405) to WS. AATW is currently supported by an Endeavour Postdoctoral Fellowship awarded by the Australian Department of Education and Training (Grant Agreement: 6534_2018) and a Marie Sklodowska-Curie Global Fellowship awarded by the Research Executive Agency of the European Commission (Grant Agreement: 797326).

**Author contributions**: B.E. and W.S. conceived and supervised the study; all co-authors conducted the fieldwork; A.A.-T.W. and J.R. conducted the molecular laboratory work; J.R. generated and analysed the morphometric data; K.S. and B.E. designed, conducted and analysed the mate-choice experiments; A.A.-T.W. analysed the genomic data; A.A.-T.W. and W.S. drafted the manuscript, with feedback from all co-authors.

**Competing interests**: The authors declare no competing interests.

**Data and materials availability**: The raw sequence reads were deposited on SRA and are available with accession numbers SRP156808 (*A. burtoni, C. horei*, and *H. stappersii*) and SRP148476 (*P. philander*). All data needed to evaluate the conclusions in the paper are present in the paper and/or the Supplementary Materials.

## Supplementary Materials

**Fig. S1.**
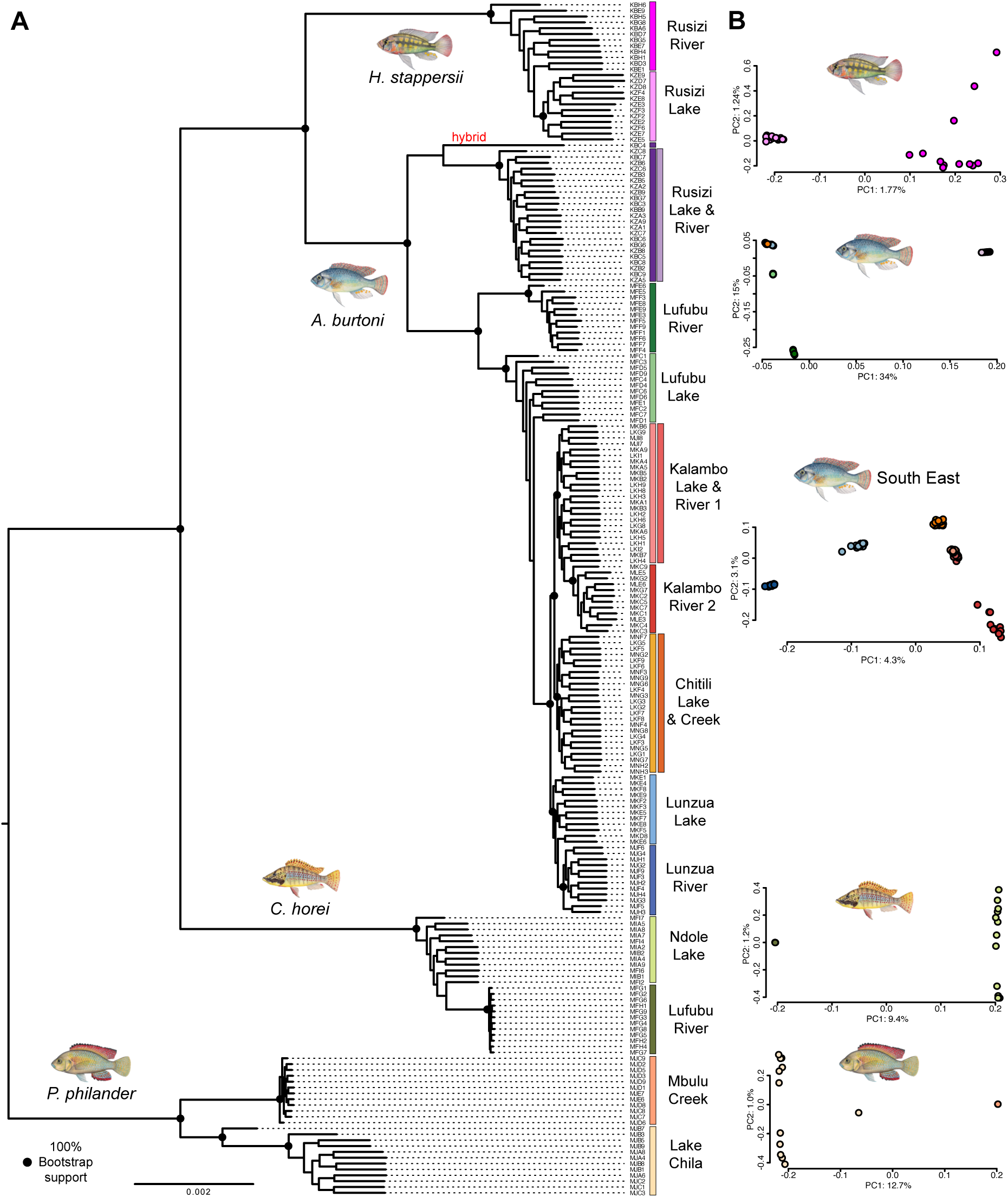
Phylogeny and genetic structure of *A. burtoni* (N=132), *H. stappersii* (N=24)*, C. horei* (N=24) and *P. philander* (N=24) (A) RAxML phylogenetic reconstruction based on a concatenated whole-genome dataset. A deep split was found between the northern and the southern populations of *A. burtoni*, confirming the findings of a recent phylogeographic study^29.^ KBC4, a putative hybrid individual excluded from the analyses, is highlighted in red. The riverine populations of *C. horei* and *P. philander* display extremely low levels of genetic diversity, suggesting that these populations are currently experiencing or have experienced a strong genetic bottleneck. (**B**) First and second axes of whole-genome principal component analyses (PCA), split per species.

**Fig. S2.**
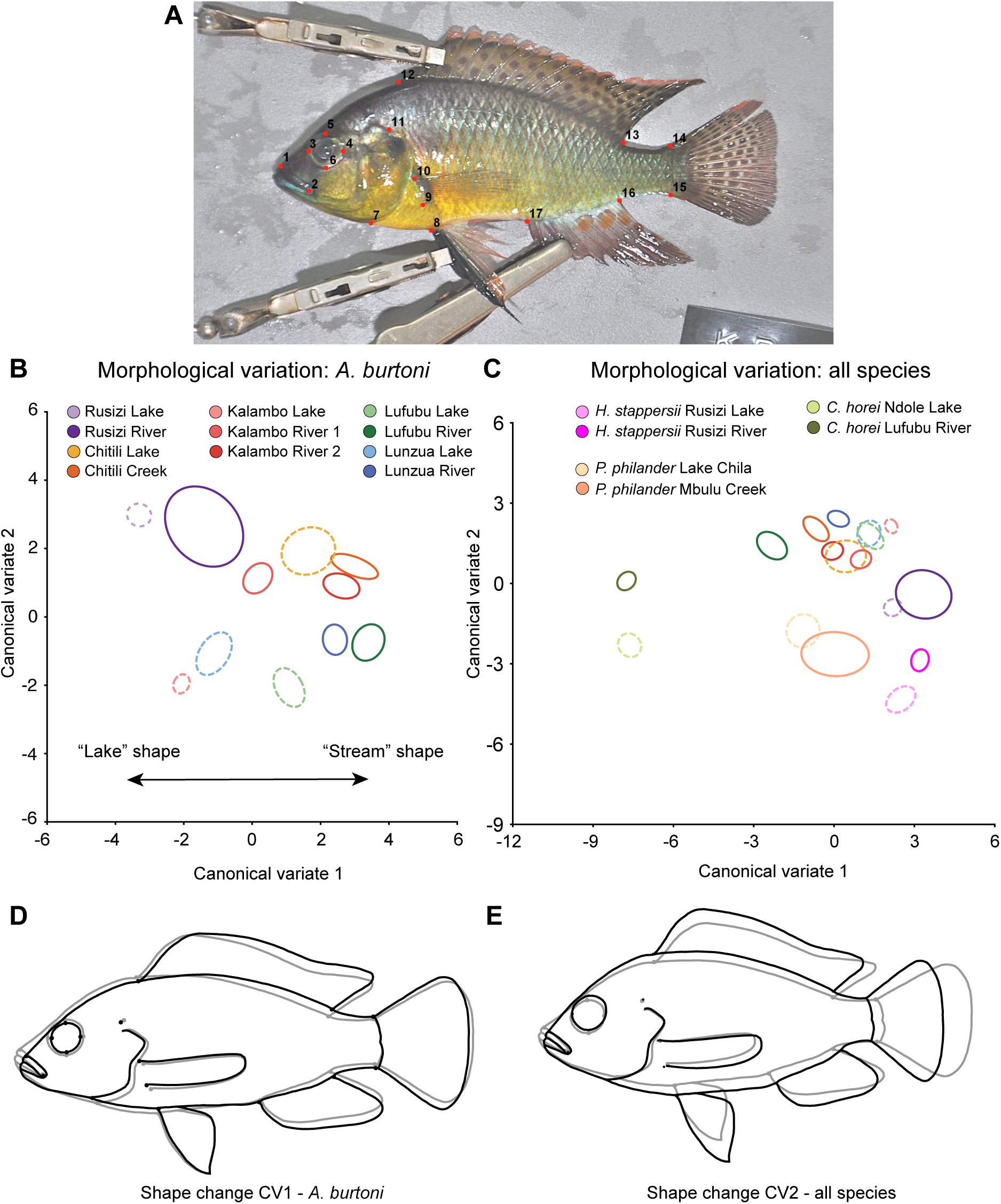
**Geometric morphometric analyses of *A. burtoni* (N=289), *H. stappersii* (N=81)*, C. horei* (N=67) and *philander* (N=31).** (**A**) Position of 17 landmarks used for geometric morphometrics. (**B**) Canonical variate analysis (CVA) of body shape for *A. burtoni* only. Lake populations outlines are shown in dashed lines. **(C)** Canonical variate analysis (CVA) of body shape for all species. Lake populations outlines are shown in dashed lines. (**D**) CV1 shape change for *A. burtoni* only (scaling factor: 10; outlines are for illustration purposes only, from light grey to dark outlines with increasing values). (**E**) CV2 shape change for all species (scaling factor: 10; outlines are for illustration purposes only, from light grey to dark outlines with increasing values).

**Fig. S3.**
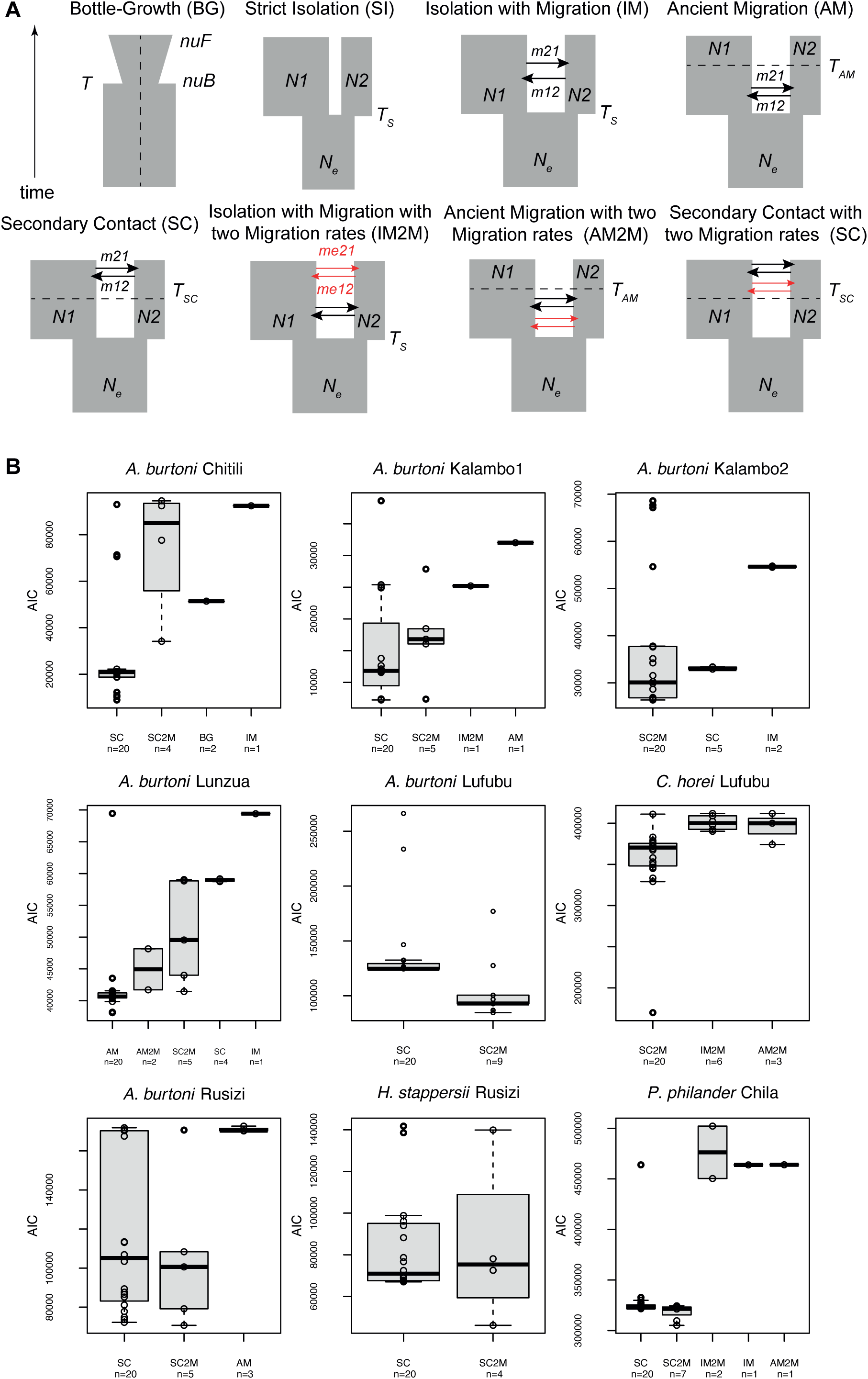
Models and results of demographic modelling based on ∂a∂i. (**A**) Schematic representation of the eight demographic models tested. Bottle-Growth (BG) model: T, Time of population size change; nuB, population size at the time of the change; nuF, current population size. Strict Isolation (SI) model: Ts, splitting time; Ne, ancestral population size; N1, current size of population 1; N2, current size of population 2. Isolation with Migration (IM) model: m21, migration rate from population 1 to population 2; m12, migration rate from population 2 to population 1. Ancient Migration (AM) model: T_AM_, Time at which migration stopped. Secondary Contact (SC) model: TSC, Time of the secondary contact. Isolation with Migration with two Migration rates (IM2M): me21, second migration rate from population 1 to population 2. me12, second migration rate from population 2 to population 1. (**B**) Most appropriate demographic model for each system in 24-32 replicates. A low AIC represents a good fit between a model and the data.

**Fig. S4.**
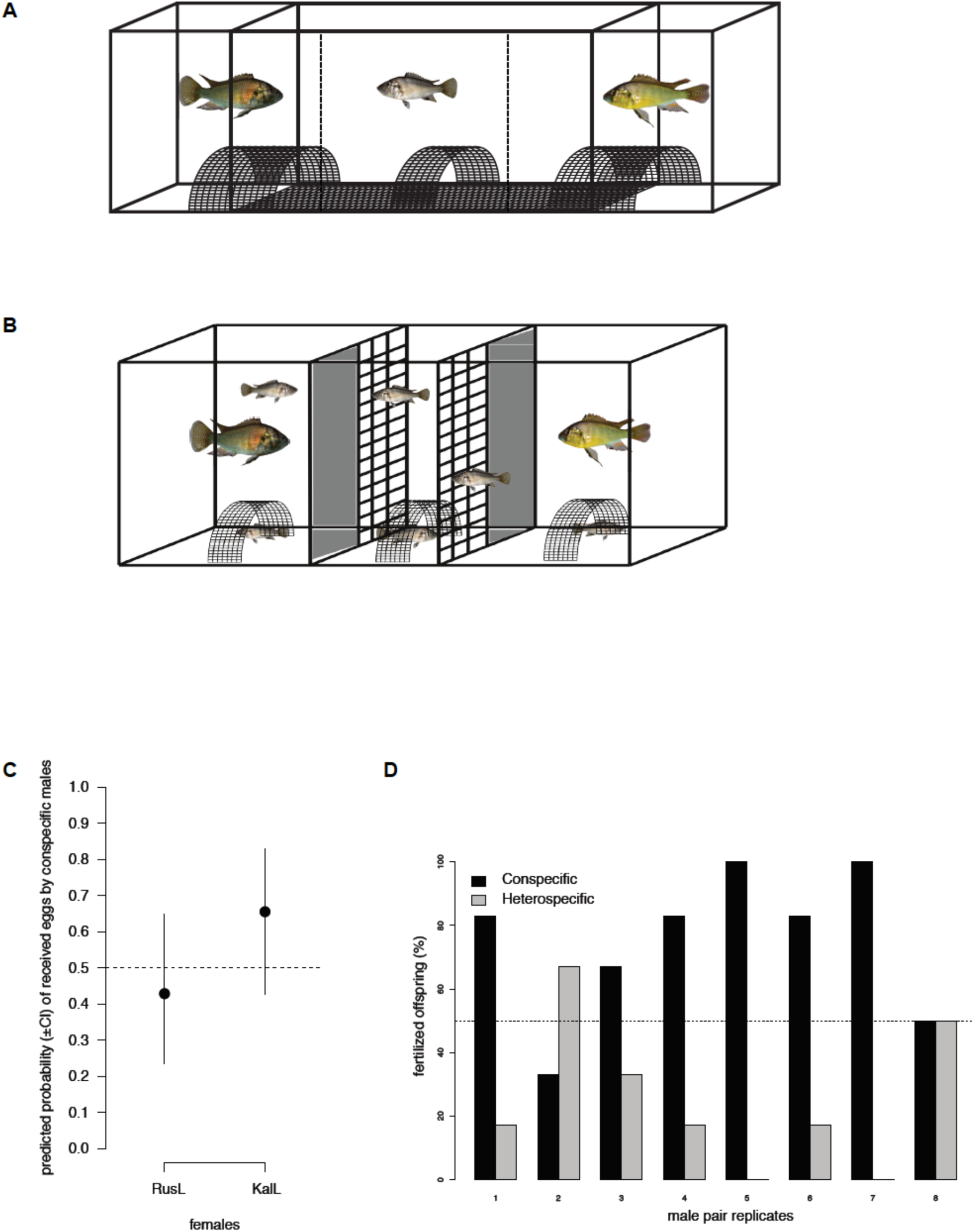
Set-up and results of mate-choice experiments. (**A**) Set-up of mate-choice experiment 1, a two-way female choice experiment based on visual cues only. A female is placed into the central aquarium equipped with an egg-trap; two stimulus males are placed in the flanking aquaria. The dashed line indicates the choice zone. (**B**) Set-up for mate-choice experiment 2, a two-way female-choice experiment with direct contact between territorial males (in the outer compartments) and the freely swimming females. (**C**) Results from the mate-choice experiment 1 (model 1). The probability of females from both populations laying eggs with their conspecific male is not different from random. RusL: Rusizi lake; KalL: Kalambo lake (**D**) Results from the mate-choice experiment 2 showing the percentage of fertilized offspring for each replicate. The females mated significantly more often with a conspecific male. Additional details on methods and results of mate-choice experiments are available in Appendix 1.

**Fig. S5.**
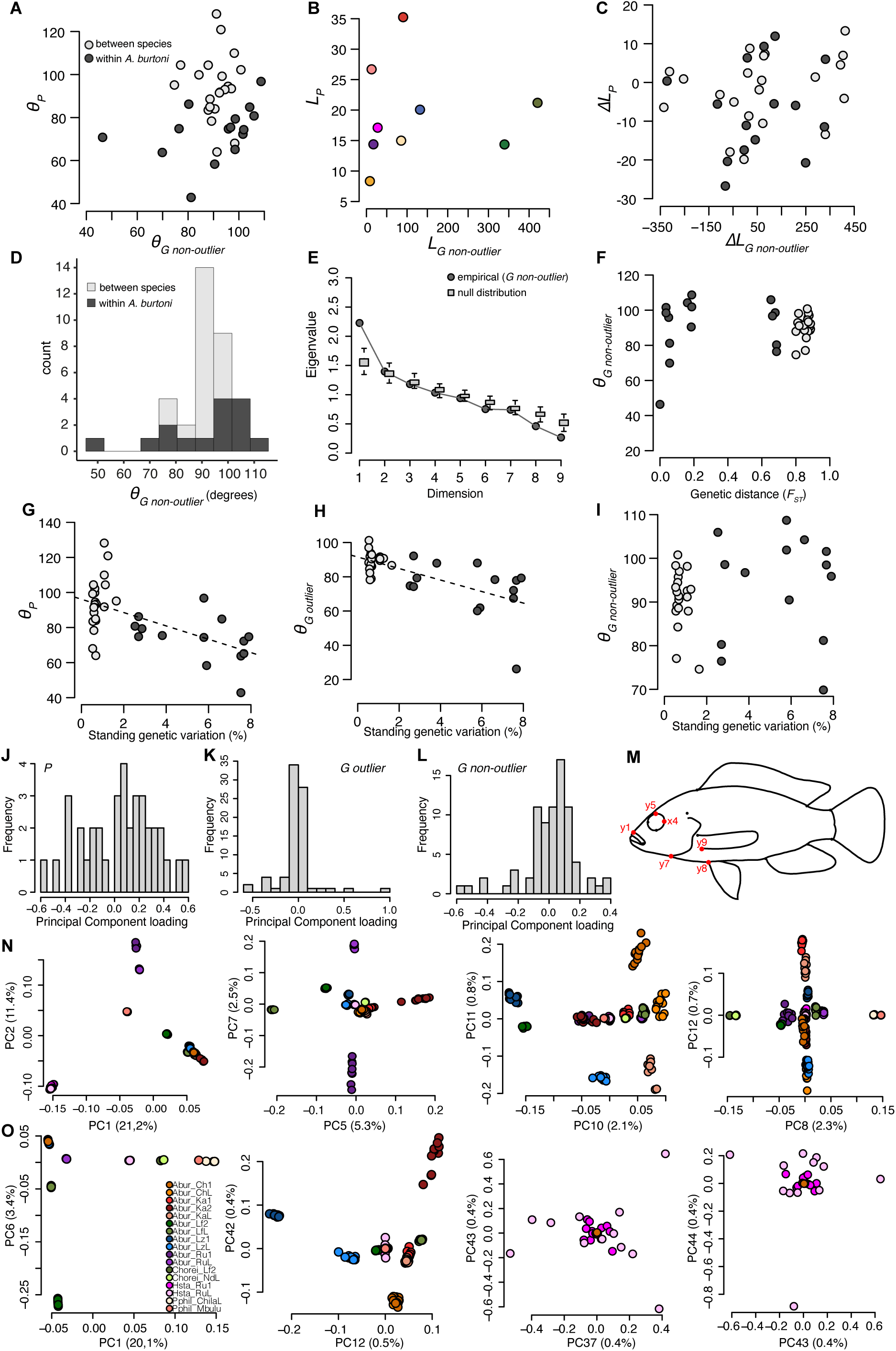
Vector analyses for genetic non-outlier data and traits underlying multivariate parallelism. (**A**) The angles of phenotypic (*_θ_P*) and genetic non-outlier (*θ_G_*) lake-stream divergence vectors are not correlated (Mantel test: P = 0.34). (**B**) The lengths of phenotypic (*L_P_*) and genetic non-outlier (*L_G_*) lake-stream divergence vectors are not correlated (Linear regression model: P = 0.93). The colour scheme is the same as in Figure 1. (**C**) The differences between phenotypic (*ΔL_P_*) and genetic non-outlier (*ΔL_G_*) vector length are not correlated (Mantel test: P = 0.35). (**D**) Histogram of the 36 (pairwise) angles between lake-stream non-outlier genetic divergence vectors (*θ_G_*) in degrees. (**E**) a multivariate analysis of genetic (non-outlier) parallelism reveals one significant dimension of parallel evolution (empirical first eigenvalue is higher than the null Wishart distribution). **(F**) The angles of genetic non-outlier divergence vectors (*θ_G_*) and genetic (*F_ST_*) distances between lake populations are not correlated (Mantel test: P = 0.48). (**G**) The angles of phenotypic divergence vectors (*_θ_P*) and the amount of standing genetic variation (SGV) between lake populations are negatively correlated (Mantel test P = 0.0061; R^2^ = 0.32). **(H**) The angles of genetic outlier divergence vectors (*θ_G outlier_*) and the amount of standing genetic variation (SGV) between lake populations are negatively correlated (Mantel test P=0.0117; R^2^ = 0.43). **(I**) The angles of genetic non-outlier divergence vectors (*θ_G_*) and the amount of standing genetic variation (SGV) between lake populations are not correlated (Mantel test P = 0.19). (**J**) Histogram of principal component loadings for each phenotypic trait from the first principal component. (**K**) Histogram of principal component loadings for each genetic outlier trait from the first principal component. (**L**) Histogram of principal component loadings for each genetic non-outlier trait from the first principal component. (**M**) Six landmark coordinates with the largest loading values from panel J are highlighted. (**N**) Eight genomic (outlier) principal components with the largest loading values from panel K are highlighted. Legend on panel O. (**O**) Seven genomic (non-outlier) principal components with the largest loading values from panel L are highlighted.

**Fig. S6.**
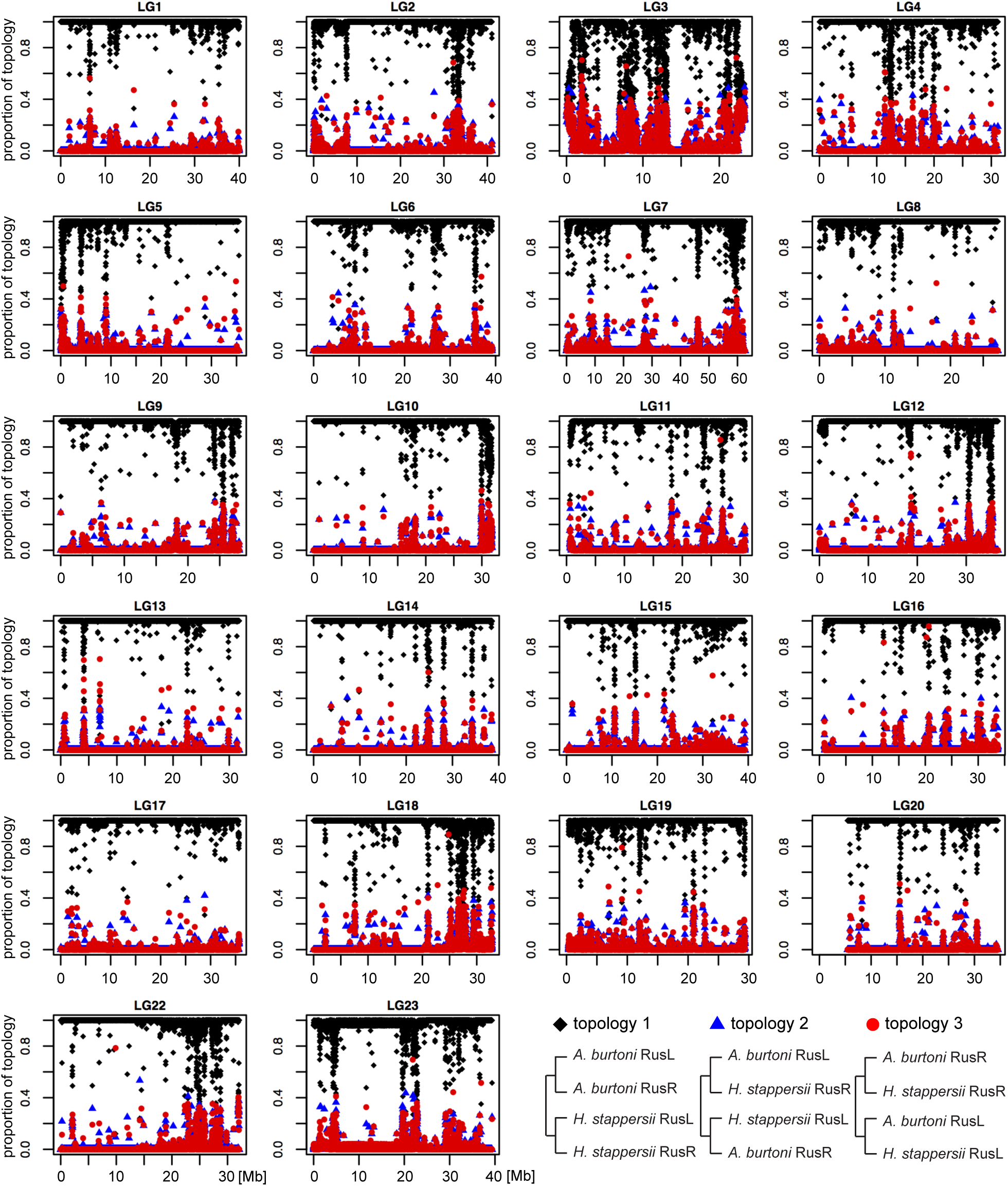
Topology weighting results per linkage group from *A. burtoni* and *H. stappersii* from the Rusizi system. Topology weighting analysis (*Twisst*) reconstructed fixed-length 5 kb windows phylogenies. The species topology (topology 1) is recovered in all cases and highlights that no introgression was detected between *A. burtoni* and *H. stappersii* in sympatry. RusL: Rusizi lake. RusR: Rusizi River.

**Fig. S7.**
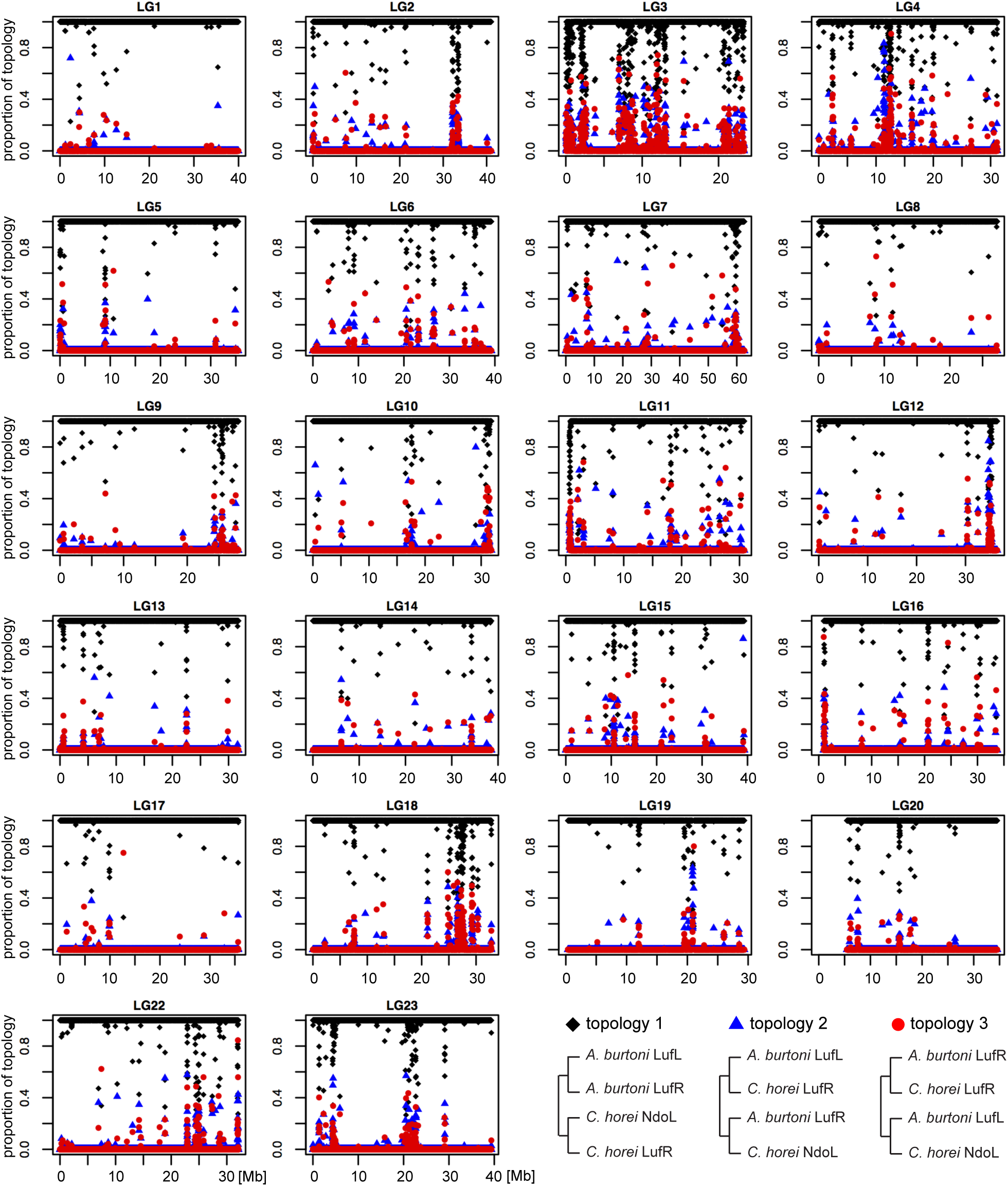
Topology weighting results per linkage group from *A. burtoni* and *C. horei* from the Lufubu system. Topology weighting analysis (*Twisst*) reconstructed fixed-length 5 kb windows phylogenies. The species topology (topology 1) is recovered in all cases and highlights that no introgression was detected between *A. burtoni* and *C. horei* in sympatry. LufL: Lufubu lake. LufR: Lufubu River. NdoL: Ndole lake.

## Supplementary tables (in a separate excel file)

**Table S1**: Individual measurements and genome statistics.

**Table S2**: Details on sampling localities, sample sizes and genome-wide population statistics.

**Table S3**: Pairwise body shape differentiation among all populations: Procrustes (upper triangular matrix) and Mahalanobis (lower triangular matrix) distances from the CVA (Fig. S2). Significant body shape differences (P< 0.05) are highlighted in bold.

**Table S4**: Population parameters inferred with ∂a∂i simulations.

**Table S5**: Number and localization of differentiation regions per system and per linkage group.

**Table S6**: List of 637 outlier candidate genes from the differentiation regions of each system and their respective GO annotations.

**Table S7**: List of 25 outlier candidate genes found in the overlap of differentiation regions among systems.

**Table S8**: List of 367 outlier candidate genes from the overlap of the three outlier core sets from *A. burtoni* northern populations, *A. burtoni* southern populations and *H. stappersii* populations.

**Table S9**: Convergence/divergence in body shape among lake-stream population pairs. Positive values indicate divergence. Negative values indicate convergence (in bold). Sympatric population pairs are highlighted in red.

**Appendix 1: Detailed methods and results of the mate-choice experiments. (in a separate pdf file)**

## Notes

### Competing Interest Statement

The authors have declared no competing interest.

