## Appendix 1 for "Diversification dynamics and (non-)parallel evolution along an ecological gradient in African cichlid fishes"

### Appendix 1: Detailed methods and results of the mate-choice experiments

#### Supplementary methods

We tested for reproductive isolation between two geographically distant and genetically divergent populations: Rusizi Lake and Kalambo Lake. Fish from the south of Lake Tanaganyika, Zambia were imported in 2011 and from the northern part of Lake Tanaganyika, near Bujumbura, Burundi in 2014. Males from each population were kept in separate aquaria (100x50x50 cm), in individual mesh cylinders provided with a half clay pot as a territorial hideout. Females from each population were kept separately in female only tanks (100x50x50 cm). Aquaria were maintained at standardized conditions (24°C and a 12:12 h light:dark cycle). Fish were fed twice a day with flake food and once a day with Artemia. All laboratory mate choice experiments were performed at the Zoological Institute of the University of Basel under the permission of the Cantonal Veterinary Office, Basel, Switzerland (permit numbers: 2536, 26037).

*Experiment 1: Visual cues only*

A two-way female choice set up (*1*, *2*) was used to test whether females preferred males of their own population when only visual cues are available (Fig. S4A). In each experimental round, we placed a gravid female of either population (n = 44) in a central tank (60 x 30 x 30 cm) and allowed visual contact with two males from Rusizi Lake and Kalambo Lake presented in two outer tanks (40 x 25 x 25 cm). The paired males were size matched in standard body length (SL) as precisely as possible (number of male pairs = 8; mean SL difference ± standard deviation [SD] = 0.51 ± 0.82 mm; range 0.0 – 3.41 mm) and introduced at least 24 hours before the start of each experimental round to allow for acclimation and territorial behaviour to develop. In each experimental round, the female was able to see and interact with both males of the stimulus pair and laid eggs within a period of few hours up to 12 days. The experiment was terminated if the female did not lay eggs within this time period. Because of the grid placed in the aquaria, eggs laid by the female would fall into this "egg-trap" before the females were able to take them into their mouth for incubation. The "egg-trap", which completely covered the floor of the female tank, made it possible to assess if the female laid the eggs in front of the Rusizi Lake male, the Kalambo Lake male, or in front of both. The position of the laid eggs was used as a measure for female preference.

After the introduction, female behaviour was recorded using a Sony handicam (HDR-X550VE, 12.0 mega pixels) for one hour and male behaviour was recorded for the same amount of time with GoPro cameras (HERO3 Silver Edition HD3.02.03.00; one per outer tank). The videos were later analysed with QuickTime player (v. 10.0) to assess the amount of time a female spent, and the time of direct activity (interaction time), with each male. For males, the time of activity, which we defined as the time males were not resting or hiding, the frequency of different behaviours i.e. lateral display, quivering, charge, facial bar, clicking (*3*–*5*) and the coloration (yellow or blue) were recorded. The side allocated to the male of each population was swapped after every second experimental run to control for directional bias; and females were tested randomly regarding the source population. Because many of the females (89%) laid their eggs exclusively next to one of the males, the data was coded into 1 and 0 to circumvent the problem of zero inflation in statistical analyses. The data was coded as 1 if the conspecific male received more than 50% of the eggs and as 0 if the conspecific male received fewer than 50% of the eggs (there was no case were both males received exactly 50% of the eggs).

The binomial data were then analysed with generalized linear mixed models (GLMMs) with a logistic link function using the package lme4 (*6*) in R (*7*) (v. 3.1.3). To correct for multiple testing of male pairs, pairs were used as a random factor. Three different models were used: model 1 tested whether the probability of the females spawning with the conspecific male was significantly different from 0.5, which was indicated by an intercept on the logit scale different from 0. Female spawning decisions were used as a response variable. Model 2 tested first (1) if females a) spent more time, or b) interacted more with the conspecific male relative to the heterospecific male; second (2) if females a) spent more time, or b) interacted more with the male they later chose in comparison to the rejected male, and third (3) if females a) spent more time, or b) interacted more with the more active male. An observation level was included as a random effect to account for the extra variance in the data. Model 3 tested if the final choice of a female depended on a) the time females spent with a respective male, b) the interaction time of females with a respective male, c) the time males were active, d) the frequency of behaviours of males (pooled and separately), e) coloration of males. We further analysed if there were differences in activity of males between populations, or between chosen and non-chosen males (as in Model 2).

*Experiment 2: direct contact*

The partial partition method (*8*) was used to infer female spawning decisions in a multi-sensory setting with free contact between females and males. A single tank (150x50x50 cm) was subdivided into 3 equally sized compartments (30x50x50 cm) by plastic grids (Fig. S4B). The middle compartment offered a resting and hiding place for the females whereas the two outer compartments served as male territories. The grid size was chosen to allow the smaller females to migrate between the three compartments, and to prevent direct contact between the larger males to exclude male-male competition. Additionally, we positioned opaque plates on each half of the grid to prevent visual contact between males. We conducted eight trials, each time using two males and six females (number of males = 16; number of females = 46; in one replicate only 4 females were available due to sample size restrictions). In every trial two males, one of each population, were placed in the opposite compartments, where both males were size matched and acclimation time was at least 24 hours, as in experiment 1. Then a total of 4-6 females (2-3 females of each population) were introduced in the middle compartment. After each time two females had spawned, we switched the position of the males to avoid compartment effects.

Mouthbrooding females were caught and the fry was removed and anesthetized using clove oil before being transferred into ethanol for later DNA extraction. A fin clip of the caudal fin of the female was taken, and after an experimental round was terminated (all females spawned and were removed) a fin clip of the potential males was also taken for later DNA extraction. We recorded fertilization rate by counting the number of fertilized vs. unfertilized eggs. DNA of ten larvae per clutch, their corresponding mothers and the putative fathers was used for paternity testing with microsatellite markers Ppun5, Ppun7, Ppun21 (*9*), UNH130 (*10*) and Abur82 (*11*). The amplified DNA samples were genotyped on an Applied

Biosystems (ABI) 3130xl genetic analyser and sized in comparison to LIZ 500(-250) (ABI) internal size standard. The genotypes were determined manually with the software Peakscanner (v. 1.0) and paternity was determined using the software Cervus (*12*) (v.3.0) with no mismatch allowed. For statistical analyses we coded the data as in experiment 1 and analysed each of the eight replicates separately, and then pooled together, using binomial tests with a probability of 0.5 and a confidence interval of 0.95, to check if the females spawned more with the conspecific males.

### Supplementary results

*Experiment 1: Visual cues only*

In our female two-way choice experiment pairs of size-matched males, one from Rusizi Lake and one from Kalambo Lake, were presented to a focal female in two outer tanks arranged on both sides of the central female tank (Fig. S4A). Out of 44 females, 24 females laid more than 50% of the eggs with the conspecific male (11 Rusizi Lake females, 13 Kalambo Lake females). Males from neither population were more likely to receive more eggs from females (model 1; GLMM, number of females = 44, number of male pairs = 8, $z = 0.06$, $p = 0.55$). Disentangling the results for the two populations also indicated random choice for both Rusizi and Kalambo Lake females (GLMM, number of male pairs = 8; number of females Rusizi Lake = 21, z Rusizi Lake = -0.63, p Rusizi Lake = 0.53; number of females Kalambo Lake = 23, z Kalambo Lake = 1.34, p Kalambo Lake = 0.18; Fig. S4C).

For model 2 we used the time a female spent with either male within the first 30 minutes of the experiment. During this time, females did not spend more time with the conspecific nor with the heterospecific male when analysed pooled or separately (model 2, (1a) GLMM, number of females = 44, number of male pairs = 8, $z = 0.84$, $p = 0.40$). Females did not interact more with neither the conspecific nor with the heterospecific male (model 2, (1b) GLMM, number of females = 44, number of male pairs = 8, $z = 1.24$, $p = 0.22$). Females showed a trend towards interacting more with the male they later spawned with (model 2 (2b) GLMM, number of females = 44, number of male pairs = 8, $z = 1.68$, $p = 0.09$) and females of the Kalambo Lake population interacted more with the chosen male (GLMM, number of females = 23, number of male pairs = 8, $z = 2.604$, $p = 0.009$). Nevertheless, there was no significant interaction between the choice of egg-laying and interaction time of females with the chosen male, as shown by the results of Model 3b (GLMM, $z = 0.998$, $p = 0.32$). Analyses (3) of Model 2, with all females pooled (GLMM, number of females = 44, number of male pairs = 8, $z = 1.94$, $p = 0.05$) suggested that females tended to interact more with males that were more active. When females were analysed separately for each population, Kalambo Lake females interacted more with males that were more active (GLMM, number of females = 23, number of male pairs = 8, $z= 3.09$, $p = 0.002$).

The results from model 3 showed that Rusizi Lake females tended to lay their eggs more in front of more active males (GLMM, $z = 1.47$, $p = 0.08$); and tended to spawn more often with yellow males (GLMM, $z = 1.77$, $p = 0.08$). Kalambo Lake females tended to prefer to lay their eggs with males that exhibit a higher frequency of lateral display (GLLM, $z = 1.72$, $p = 0.08$). Kalambo Lake males tended to be more active compared to Rusizi Lake males during the experimental runs with Rusizi Lake females (GLMM, $z = 1.83$, $p = 0.07$), and with Kalambo Lake females (GLMM, $z = 1.657$, $p = 0.09$). Rusizi Lake males showed yellow coloration more often in comparison to Kalambo Lake males ($z = 1.87$, $p = 0.06$). When testing for differences in activity between chosen and non-chosen males, we found that the males chosen by Kalambo Lake females tended to have higher frequencies in overall behaviours (GLMM, $z = 1.97$, $p = 0.05$) and higher frequencies in lateral display (GLMM, $z = 2.165$, $p = 0.03$).

*Experiment 2: direct contact*

In the partial partition experimental set up with multiple cues available, females mated significantly more often with a conspecific male (binomial test: $n = 46$, $p < 0.001$, Fig. S4D). This result was consistent when each population was analysed separately (binomial test Rusizi Lake: $n = 23$, $p = 0.01$; Binomial test Kalambo Lake: $n = 23$, $p = 0.03$). In six out of eight replicates, the probabilities of females to spawn with their conspecific male ranged between 0.66 and 1; in one replicate the probability to spawn with the conspecific male was 0.33, and in the replicate where only four females were tested the probability to spawn with the conspecific male was 0.5. Fertilization rate was 100% and each brood was fertilized by a single male. Due to the fact that size matched male pairs were tested with several (4-6) females, we additionally applied a general linear mixed model with pairs as a random factor to control for multiple testing. The random effects were not significant and the results of this test were consistent with the previous ones (GLMM, number of females = 46, number of male pairs = 8, $z = 2.617$, $p = 0.008$; Rusizi Lake: number of females = 23, $z = 2.534$, $p = 0.01$; Kalambo Lake: number of females = 23, $z = 2.212$, $p = 0.02$).
